# Unexpected post-glacial colonisation route explains the white colour of barn owls (*Tyto alba*) from the British Isles

**DOI:** 10.1101/2021.04.23.441058

**Authors:** Ana Paula Machado, Tristan Cumer, Christian Iseli, Emmanuel Beaudoing, Anne-Lyse Ducrest, Melanie Dupasquier, Nicolas Guex, Klaus Dichmann, Rui Lourenço, John Lusby, Hans-Dieter Martens, Laure Prévost, David Ramsden, Alexandre Roulin, Jérôme Goudet

## Abstract

The climate fluctuations of the Quaternary shaped the movement of species in and out of glacial refugia. In Europe, the majority of species followed one of the described traditional postglacial recolonization routes from the southern peninsulas towards the north. Like most organisms, barn owls are assumed to have colonized the British Isles by crossing over Doggerland, a land bridge that connected Britain to northern Europe. However, while they are dark rufous in northern Europe, barn owls in the British Isles are conspicuously white, a contrast that could suggest selective forces are at play on the islands. However, analysis of known candidate genes involved in colouration found no signature of selection. Instead, using whole genome sequences and species distribution modelling, we found that owls colonised the British Isles soon after the last glaciation, directly from a white coloured refugium in the Iberian Peninsula, before colonising northern Europe. They would have followed a yet unknown post-glacial colonization route to the Isles over a westwards path of suitable habitat in now submerged land in the Bay of Biscay, thus not crossing Doggerland. As such, they inherited the white colour of their Iberian founders and maintained it through low gene flow with the mainland that prevents the import of rufous alleles. Thus, we contend that neutral processes likely explain this contrasting white colour compared to continental owls. With the barn owl being a top predator, we expect future research will show this unanticipated route was used by other species from its paleo community.

## Introduction

The dramatic climate fluctuations of the Quaternary were key in shaping the global distribution of species and communities observed today (Ficetola, Mazel, & Thuiller, 2017; Hewitt, 2000). During the last glaciation, northern Europe was largely covered by ice caps, and the resulting lower sea levels unveiled an expanded coastline widely different from that of today. The inhospitable conditions throughout the continent forced many temperate species into warmer refugia, most commonly the southern peninsulas of Iberia, Italy and Balkans (Hewitt, 1999, 2011). Once temperatures started increasing about 18 thousand years BP, ice sheets melted, the sea rose and these species re-expanded northwards into central and northern Europe, a key step in determining their modern distribution and genetic structure across the continent. Early comparative phylogeography studies described differences in the route and timing of colonisation from each refuge population and identified the main post-glacial recolonization patterns from the south (Hewitt, 1999, 2000; Taberlet, Fumagalli, Wust-Saucy, & Cosson, 1998). However, advances in sequencing technology and the consequent increase in studies with high representation molecular markers have since provided numerous examples of alternative routes and cryptic refugia for different taxa in mainland Europe as well as on islands (Bilton et al., 1998; Deffontaine et al., 2005; García-Vázquez, Pinto Llona, & Grandal-d’Anglade, 2019; Herman et al., 2017; Stewart & Lister, 2001).

The colonisation of the British Isles by terrestrial organisms has often been described in the context of the main phylogeographic patterns, with mainland north-western Europe as its origin (Hewitt, 1999, 2000; Montgomery, Provan, McCabe, & Yalden, 2014). Such a route would have been facilitated by Doggerland, a large land bridge of alluvial plains that connected Great Britain (GB) to mainland northern Europe before submerging under the north Sea 8’000 years BP (Coles, 1998; Ward, Larcombe, & Lillie, 2006). Most terrestrial vertebrates of GB do appear to have arrived via Doggerland, as evidenced by the similarity between its mammal fauna and that of northern rather than southern Europe (Montgomery et al., 2014). Nonetheless, some species believed to have followed this path were found to have had glacial refugia on the islands themselves (Stewart & Lister, 2001), including plants (Kelly, Charman, & Newnham, 2010), amphibians (Snell, Tetteh, & Evans, 2005; Teacher, Garner, & Nichols, 2009) and mammals (Boston, Ian Montgomery, Hynes, & Prodöhl, 2015; Lister, 1984). Some taxa revealed other surprising post-glacial patterns such as colonization of the British Isles from multiple refugia in independent waves (badger: O’meara et al. 2012; water vole: Brace et al. 2016) and even separate colonisation of Ireland and GB (stoat: Martínková et al. 2007).

Barn owls (*Tyto alba*) recolonised western Europe following the last glaciation from a refugium in the Iberian Peninsula (Antoniazza et al., 2014; Burri et al., 2016). On the mainland, barn owl ventral plumage colouration follows a latitudinal cline ranging from mostly white in the southern populations to dark rufous in the north (Antoniazza, Burri, Fumagalli, Goudet, & Roulin, 2010; Antoniazza et al., 2014). Despite their post-glacial expansion route, the clinal variation in colour was not a neutral by-product of range expansion, but was rather created and maintained by an independent post-glacial selective process (Antoniazza et al., 2014). The genetic basis of this pheomelanin-based trait is not fully understood, but a specific non-synonymous variant (V126I) in the melanocortin-1 receptor (*MC1R*) gene has been found to explain roughly 30% of its variation in Europe (San-Jose et al., 2015). The derived *MC1R* rufous allele produces the darkest owl phenotypes and follows the European colour cline of increasing frequency with latitude (Burri et al., 2016).

It is hypothesised that, given their aversion to crossing large water bodies, barn owls recolonized Great Britain following the traditional route by crossing over Doggerland (Martin, 2017). However, barn owls from the British Isles are famously white (Martin, 2017; Roulin & Randin, 2016) in stark contrast to their darker mainland counterparts at similar latitudes. Over-land expansion from a north-western European population, inhabited mostly by rufous owls with 10% - 45% rufous *MC1R* allele, would be at odds with the whiteness of the GB population. This disparity is especially startling, given that rufous individuals disperse further than white ones (Roulin, 2013; van den Brink, Dreiss, & Roulin, 2012), and would thus be more likely to colonise the islands in the first place. Finally, with GB being a recently isolated island, its avifauna is very similar to that of continental Europe (albeit less species rich), and examples of such phenotypic divergence from the mainland are rare; the barn owl is thus an intriguing exception. Being sensitive to extreme cold (Altwegg, Roulin, Kestenholz, & Jenni, 2006), a northern refugium seems unlikely. However, such phenotypic disparity suggests that, unless strong selective pressure is involved, the colonisation timing and route of barn owls of the British Isles might have been less straightforward than has been assumed.

Here, we address the post-glacial colonisation history of barn owls in the British Isles in light of the puzzling whiteness of their plumage. First, with a new broad sampling of 147 individuals from western Europe, we confirm that owls from the British Isles do not fit into the expected colouration and *MC1R* pattern of the mainland, with darker individuals at higher latitudes. Taking advantage of a highly contiguous newly-assembled reference genome and using the whole-genome sequences of 61 individuals, we use the neutral genetic structure to model the demographic history of barn owl colonisation of the northern part of Europe and the British Isles from a glacial refugium in Iberia. Then, we use ringing data to support estimations of current gene flow. Lastly, we investigate the potential role of other colour-linked genes in maintaining the phenotypic disparity in plumage colour between the British Isles and mainland Europe.

## Materials & Methods

### Tissue sampling, *MC1R* genotyping and colour measurement

In total, 147 individual barn owls were sampled for this study from six European populations (Sup. Table 1): Ireland (IR), Great Britain (GB), France (FR), Switzerland (CH), Denmark (DK) and Portugal (PT). A denser sampling was performed in the British Isles (n=113) as this was the first time these populations were studied, while for the mainland populations data was already available (Burri et al., 2016). Genomic DNA was extracted from blood, feathers or soft tissue using the DNeasy Blood & Tissue kit (Qiagen, Hilden, Germany) following the manufacturer’s instructions, including RNA digestion with RNase A. A previously established allelic discrimination assay (San-Jose et al., 2015) was used to molecularly determine individual genotypes at the amino acid position 126 of the Melanocortin 1 receptor (*MC1R*) gene of the 147 individuals (Sup. Table 1). Additional allelic frequencies at this locus published in Burri *et al.* (2016) from the mainland populations of interest to this study were used for context (N=247 individuals; Appendix 1).

For all individuals with available breast feathers (*n*=145), pheomelanin-based colour was estimated as the brown chroma of the reflectance spectra (for detailed description see Antoniazza *et al*. 2010). Briefly, the brown chroma represents the ratio of the red part of the spectrum (600–700 nm) to the complete visible spectrum (300–700 nm). The reflectance of four points of the top of three overlapping breast feathers was measured using a S2000 spectrophotometer (Ocean Optics, Dunedin, FL) and a dual deuterium and halogen 2000 light source (Mikropackan, Mikropack, Ostfildern, Germany). An individual’s brown chroma score was obtained as the average of these four points. Brown chroma data from Burri *et al.* (2016) were used to complete the dataset, using the same individuals as for the *MC1R* analysis (Appendix 1). Given the marked non-normality of the data, a non-parametric Kruskal-Wallis test was performed to detect differences in coloration between the six populations. Further, a Pairwise Wilcoxon Rank Sum test was used to identify significant differences between pairs of populations using a Bonferroni correction.

### New reference genome

As the available reference genome for the European *Tyto alba* was fragmented (Ducrest et al., 2020), a new reference was produced in order to achieve a near chromosome-level assembly. A full description of the process and its detailed results are given in Appendix 2. Briefly, a long-read PacBio library was produced from a blood sample of a Swiss individual at an expected coverage of 100x for the barn owl’s 1.3Gb genome. FALCON and FALCON-Unzip v.3 (Chin et al., 2016) were used to assemble PacBio reads. Then, a high molecular weight DNA Bionano optical mapping library was used to assemble PacBio contigs into scaffolds. Finally, repeated regions were identified using RepeatModeler v.1.0.11 (Smit & Hubley, 2008-2015) and masked with RepeatMasker v.4.0.7 (Smit, Hubley, & Green, 2013-2015). Coding regions were identified using the Braker2 pipeline v.2.0.1 (Brůna, Hoff, Lomsadze, Stanke, & Borodovsky, 2020; Hoff, Lange, Lomsadze, Borodovsky, & Stanke, 2016; Hoff, Lomsadze, Borodovsky, & Stanke, 2019; Stanke, Diekhans, Baertsch, & Haussler, 2008; Stanke, Schöffmann, Morgenstern, & Waack, 2006).

### Whole-genome resequencing and SNP calling

For the population genomics analyses of this study, the whole genomes of 61 out of the 147 individual barn owls were sequenced (Sup. Table 1). In addition, one eastern (*T. javanica* from Singapore) and one American barn owl (*T. furcata* from California, USA) were used as outgroups. See Supplementary Methods for a complete description of the library preparation, sequencing, SNP calling and filtering. Briefly, individual 100bp TruSeq DNA PCR-free libraries (Illumina) were sequenced with Illumina HiSeq 2500 high-throughput paired-end sequencing technology at the Lausanne Genomic Technologies Facility (GTF, University of Lausanne, Switzerland). The bioinformatics pipeline used to obtain analysis-ready SNPs was adapted from the Genome Analysis Toolkit (GATK) Best Practices (Van der Auwera et al., 2013) to a non-model organism following the developers’ recommendations, producing a full dataset of 6’721’999 SNP for the 61 European individuals with an average coverage of 21.1x (3.36 SD).

### Population structure and genetic diversity

To investigate population structure among our samples, sNMF v.1.2 (Frichot, Mathieu, Trouillon, Bouchard, & François, 2014) was run for K 2 to 6 in 25 replicates to infer individual clustering and admixture proportions. For this analysis, singletons were excluded and the remaining SNPs were pruned for linkage disequilibrium (LD) with PLINK v1.946 (Purcell et al., 2007; parameters - indep-pairwise 50 10 0.1) as recommended by the authors, yielding 319’801 SNP. The same dataset was used to perform a Principal Component Analysis (PCA) with the R package SNPRelate (Zheng et al., 2012). Treemix (Pickrell & Pritchard, 2012) was used to infer population splits in our data, using the LD-pruned dataset further filtered to include no missing data (180’764 SNP). To detect meaningful admixture between populations, 10 replicates were run for 0 to 8 migration events, with the tree rooted on the PT population, representative of the glacial refugium. An extra run without migration events was conducted with a north-American owl as an outgroup in the dataset to verify that the root did not affect the topology of the tree.

To estimate population statistics, individuals found to be mis-assigned to their given population based on genetic structure analyses (PCA and sNMF) were removed so as not to bias allelic frequencies (N=3 individuals from Ireland). Individual expected and observed heterozygosity and population-specific private alleles were estimated using custom R scripts for each genetic lineage identified by sNMF with K=4. To account for differences in sample sizes, private alleles were calculated by randomly sampling 9 individuals from the larger populations (GB and central Europe) 10 times in a bootstrap-fashion and estimating the mean. Individual-based relatedness (β; Weir and Goudet 2017), inbreeding coefficient for SNP data, overall and population pairwise *F*_ST_ (B.S. Weir & Cockerham, 1984) were calculated with SNPRelate.

### Gene flow and migration analyses

#### Migration surface estimate

The Estimated Effective Migration Surface (EEMS) v.0.0.9 software (Petkova, Novembre, & Stephens, 2016) was used to visualize geographic regions with higher or lower than average levels of gene flow within our dataset. The provided tool *bed2diff* was used to compute the matrix of genetic dissimilarities, from the dataset pruned for LD produced above. The free Google Maps api v.3 tool (http://www.birdtheme.org/useful/v3tool.html) was used to draw the polygon outlining the study area in western Europe. EEMS was run with 750 demes in three independent chains of 5 million MCMC iterations with a 1 million iterations burn-in. Results were checked for MCMC chain convergence visually and through the linear relation between the observed and fitted values for within- and between-demes estimates using the accompanying R package rEEMSplots v.0.0.1 (Petkova et al., 2016). The three MCMC chains were combined to produce maps of effective migration and diversity surfaces with the provided functions in rEEMSplots.

#### Treatment and analyses of capture-recapture data

In addition to genomic data, recapture data of ringed barn owls across Europe were obtained from the EURING database (obtained in March 2020; Speek et al. 2001; du Feu et al. 2016). Specifically, we estimated the frequency of crosses over open water between GB and central and western Europe, as well as between GB and Ireland. To do so, we kept records of birds that had been recaptured at least once after ringing (n=94’797 recaptures, n=80’083 individuals, from 1910 to 2019) and filtered the accuracy of the “time of capture” parameter to a period of within 6 weeks of the reported date to exclude potentially unreliable data points. We extracted the number of birds ringed and recaptured in GB and Ireland, as well as in the countries that produced or received migrant birds from these islands and central Europe (Belgium, Denmark, France, Spain, Germany, Switzerland and The Netherlands). Crosses between islands and to/from the mainland are reported and include birds that were found dead in the sea (n=8). All counts and percentages reported are relative to the number of individual birds recaptured (rather than number of recapture events, as a single bird can be recaptured multiple times).

### Post-glacial species distribution

To support the demographic scenarios tested in the following section, we modelled the past spatial distribution of barn owls in western Europe, in order to identify the regions of high habitat suitability at the last glacial maximum (LGM, 20’000 years BP). A complete description of the models can be found in Supplementary Methods.. Briefly, using Maximum Entropy Modelling (MaxEnt), a presence-only based modelling tool, we built species distribution models (SDM) based on climatic variables extracted from the WorldClim database (Hijmans, Cameron, Parra, Jones, & Jarvis, 2005) at 5 arc min resolution. Then, the output of the models was transformed into a binary map of suitability in which only cells suitable in 90% of the models are presented as such in the map. All models were then projected to the mid-Holocene (6’000 years BP) and LGM (20’000 years BP) conditions extracted from WorldClim at the same resolution as current data. For each timepoint, the results of the models were merged and transformed into a binary map as for the current data.

### Maximum-likelihood demographic inference

#### Data preparation

To discriminate between different demographic scenarios for the colonisation of the British Isles by barn owls we used the software *fastsimcoal2* (Excoffier, Dupanloup, Huerta-Sánchez, Sousa, & Foll, 2013; Excoffier & Foll, 2011). Individuals and variants in the dataset used here as input went through additional filtering steps in an attempt to ensure neutrality and homogeneity between samples (Sup. Methods). Given their similarity (Fig. 1b&c), the original populations of France, Denmark and Switzerland were combined into a central European population (EU). The remaining populations were Portugal (PT), Great Britain (GB) and Ireland (IR), with 8 individuals each (Sup. Table 1). Population pairwise SFS were produced from the filtered dataset of 739’168 SNP.

**Figure 1.**
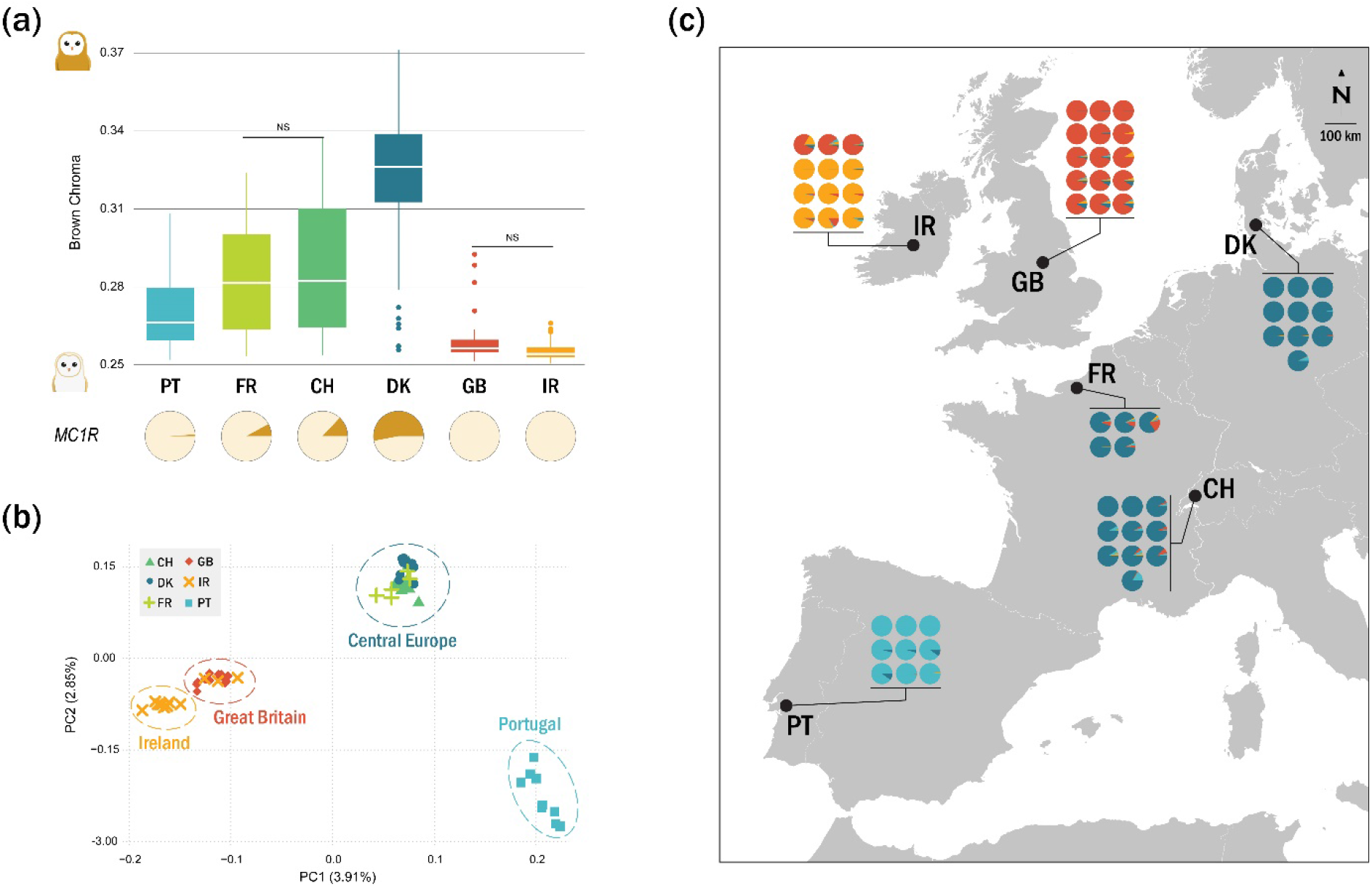
Colouration and genetic structure of barn owl populations in western Europe. (a) Brown chroma distribution and *MC1R* allelic frequencies of each studied population (total *N*=145). Higher brown chroma indicates redder owls. NS denotes the non-significant pairwise comparisons. The pies below the plot illustrate the populations’ *MC1R* allelic frequencies: the rufous allele in brown and the white in beige. (b) PCA based on the pruned SNP set of the 61 individuals whose whole genome was re-sequenced. Point shape and colour denote populations according to the legend. Dashed circles enclose sample clusters identified in sNMF. Values in parenthesis indicate the percentage of variance explained by each axis. (c) Population structure. Small pie charts denote the individual proportion of each of k=4 lineages as determined by sNMF. Black dots are located at the approximate centroid of each sampled population.

#### Demographic scenarios and parameters

Three different scenarios of colonization of central Europe and the British Isles from the Iberian Peninsula were simulated (Figure 3), distinguishable by the difference in timing and origin of the insular populations: north-western (NW) European origin, Iberian origin and insular refugium. Each scenario was further split in two versions (A and B) to accommodate small changes in topology. For all scenarios, wide search ranges for initial simulation parameters were allowed for population sizes, divergence times and migration rates while accounting for census and geological data (Sup. Table 7). Splits were preceded by instantaneous bottlenecks, in which the founding population size was drawn from a log-uniform distribution between 0.01 and 0.5 of current population sizes. All times were relative to the end of the last glaciation (18’000 years BP, rounded to 6000 generations ago), bounded between the present and the previous demographic event in the model.

In scenario NW European origin A, after an initial post-glaciation size expansion, the ancestral PT population colonized central Europe. From here, barn owls sequentially reached Great Britain and Ireland, potentially across the Doggerland land bridge. In version B, a smaller second glacial refugium is hypothesized to have existed in southern France, above the Pyrenees, as the founder of the central European population after the glaciation. In both versions, barn owls reached the British Isles from central Europe. In the Iberian origin scenarios, the insular populations originated directly from PT. Spatially, this could have taken place across now-submerged land in the Bay of Biscay, west of current-day France and north of Spain. Genetically, the insular birds would have been derived from the initial genetic pool in Iberia rather than from the subset in central Europe. Versions A and B of this scenario differ in the timing of colonization, with Europe being colonized before the islands in A and after in B. Lastly, the insular refugium scenarios hypothesize a separate and smaller glacial refugium in the south of the British Isles that would have been the origin of today’s populations on the islands. Such refugia have been described for some terrestrial organisms albeit not birds (Kelly et al., 2010; Ravinet, Harrod, Eizaguirre, & Prodöhl, 2014; Stewart & Lister, 2001; Teacher et al., 2009). Central Europe would be colonized post-glacially from PT. In version A and B of this scenario, the second glacial refugium would be part of an ancestral GB or IR population, respectively.

In summary, the NW European origin scenario reflects the shortest overland path based on current geography, whereas the remaining scenarios attempt to address the whiteness in the British Isles by avoiding shared ancestry with darker-coloured populations at different time scales, as well as the changes in the coastline during and after the last glaciation. For all scenarios, migration was allowed between neighbouring populations (Figure 3; Sup. Table 7).

#### Demographic inference

Demographic simulations and parameter inference were performed under a composite-likelihood approach based on the joint site frequency spectrum (SFS) as implemented in *fastsimcoal2* (Excoffier et al., 2013; Excoffier & Foll, 2011). For each scenario, 100 independent estimations with different initial values were run (Sup. Methods). The best-fitting scenario was determined based on Akaike’s information criterion (AIC; Akaike 1974) and confirmed through the examination of the likelihood ranges of each scenario as proposed in Kocher *et al.* (1989). For the best-fitting scenario, non-parametric bootstrapping was performed to estimate 95% confidence intervals (CI) of the inferred parameters. For each block-bootstrapped SFS, 50 independent parameter inferences were run for the best-fitting scenario (see Sup. Methods for a detailed explanation).

### Genome scans of colour-linked genes

Genome-wide scans were used to compare patterns of divergence and diversity between populations. SNPs were filtered to a minimum derived allelic frequency of 5%, and VCFtools was used to calculate nucleotide diversity (π) for each population and to estimate *F*_ST_ (B.S. Weir & Cockerham, 1984) between pairs of populations in 20kb sliding windows with 5kb steps across the whole genome. For our comparisons, Great Britain and Ireland were combined as British Isles; France and Switzerland as central Europe. Denmark was not included in the latter due to its markedly darker phenotype (Fig. 1a). The British Isles were compared to all other groups of individuals: white in Portugal, intermediate in central Europe and dark rufous in Denmark. Further, Portugal and Denmark were also compared.

In our genomic dataset, owls from the British Isles and Portugal carried the same genotypes at the *MC1R* mutation (100% V allele) despite there being considerably more variation in colour among Portuguese individuals (Fig. 1a). As such, we first investigated whether insular individuals showed particular diversity or divergence at the surrounding positions within the *MC1R* gene that could relate to their pure white colour. Since the *MC1R* gene in barn owls is particularly GC rich (San-Jose et al., 2015) and is embedded in a region with a lot of homopolymeric sequences, the sequencing in this region has a considerably lower coverage than the average of the genome. To account for this, the scaffold containing this gene was extracted from the raw SNP set and re-filtered with similar site thresholds as described above, except for allowing 25% overall missing data (instead of 5%), limiting the minimum individual DP to 5 (instead of 10) and the minimum minor allelic count to 3. VCFtools was used to calculate nucleotide diversity for each population and to estimate *F*_ST_ (B.S. Weir & Cockerham, 1984) between pairs of populations in 5kb sliding windows with 1kb steps along this scaffold.

Second, to widen our search to other colour-linked genes besides *MC1R*, we mapped 22 autosomal candidate genes (Appendix 3) onto the reference genome using Blast v.2.9.0 (Zhang, Schwartz, Wagner, & Miller, 2000). Windows including the candidate genes were plotted onto genomic scans (5kb windows with 1kb step) to check for overlap with peaks or drops in diversity and/or differentiation.

## Results

### *MC1R* genotyping and colour measurements

Plumage colour comparisons showed that the British Isles have the whitest owls of all measured European populations (Fig. 1a; *X*^2^ = 243.28, p < 0.001). Most pairwise comparisons were significantly different after correction, with the exception of between GB and IR owls, and between CH and FR. As for *MC1R* genotyping, notably no I allele was found among the 113 genotyped individuals of the British Isles indicating it is absent from these populations or at very low frequency.

### New reference genome

The new reference genome produced for European barn owl was a near chromosome level assembly, and has been deposited at DDBJ/ENA/GenBank under the accession JAEUGV000000000. Sequencing of the new reference genome’s PacBio library yielded 7.3 million long reads with a total sum length of unique single molecules of 135 Gbp (N50 > 31Kb) yielding a realized coverage of 108x. Its assembly with FALCON and FALCON-Unzip resulted in 478 primary contigs partially phased, and 1736 fully phased haplotigs which represented divergent haplotypes. Optical mapping with Bionano produced a final assembly of 70 scaffolds, slightly more than the barn owl’s karyotype of 46 chromosomes (Ducrest et al., 2020). The final assembly was 1.25 Gbp long, with an N50 of 36 Mbp and BUSCO score of 96.9% (see Appendix 2 Table 1 for full assembly metrics). In comparison, the previous reference assembly (Ducrest et al., 2020) had 21,509 scaffolds, with an N50 of 4.6 Mbp.

### Population structure and genetic diversity

Our dataset was composed of four main genetic clusters identified by individual ancestry analyses (sNMF) and PCA clustering. Individuals from Portugal (PT), Great Britain (GB) and Ireland (IR) belonged to their specific population ancestry, while individuals from France (FR), Denmark (DK) and Switzerland (CH) formed a single central European cluster (Fig. 1b,c; Sup. Fig. 3). Consistently, the first axis of the PCA opposed PT to GB & IR, as seen with sNMF K=2 (Sup. Fig. 3). The second axis clustered the central European individuals together and opposed them to PT (Fig. 1b). GB and IR segregate in both the first and second axes. Three barn owls sampled in Ireland showed a clear genetic signal of belonging to the Great Britain genetic cluster (Fig. 1b,c; Sup. Fig. 3). To avoid their interference in estimating allelic frequencies, they were omitted when estimating diversity and differentiation statistics.

Analyses of genetic drift with Treemix yielded a population tree with two branches splitting from PT. The first is a long branch of drift that divides into GB and IR, while the second, shorter branch, diversified into the three central European populations (Fig. 4a). Plotting the likelihood of runs and the standard error (SE) of each tree showed that including one migration event from PT to CH (migration edge weight = 0.27) considerably increased the fit of the tree to the data (Sup. Fig. 5).

The overall *F*_ST_ was 0.035. Population pairwise *F*_ST_ were the highest between Ireland and central Europe (Sup. Table 3). Overall, populations within central Europe showed the smallest differentiation (*F*_ST_ below 0.02) and the British Isles had the highest values in comparison to all mainland populations (Sup. Table 3). Diversity estimates showed higher levels in PT than in any other population and the British Isles had the lowest (Sup. Table 2). Individual relatedness was highest within IR, followed by GB (Sup. Fig. 4). On the opposite end, PT had the lowest within-population relatedness as well as with the other populations, consistent with its higher diversity.

### Migration and gene flow

The English Channel – including the strait of Dover and the southernmost part of the North Sea – was identified by Estimated Effective Migration Surface (EEMS) as a region with lower than average gene flow between populations (Fig. 2a). This corridor extended west to the Atlantic. Furthermore, this analysis highlighted a region of low gene flow between the British and Irish populations. It put a barrier in Ireland by separating the north from the rest of the island, effectively isolating the three individuals sampled in Ireland that genetically resemble the British and clustering them with GB.

**Figure 2.**
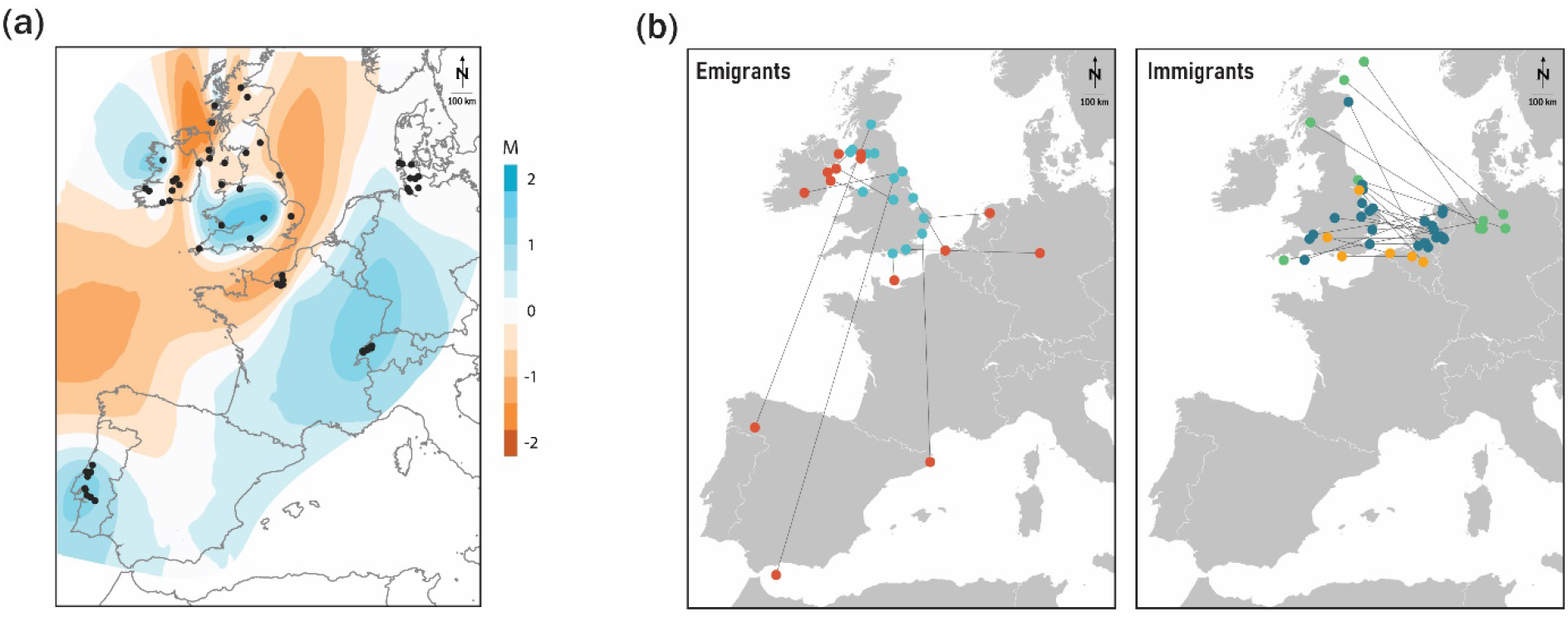
Barn owl gene flow and dispersal between the British Isles and mainland Western Europe. (a) Estimated effective migration surface (EEMS) based on whole-genome data. Blue and orange shading denote regions of higher and lower than average gene flow, respectively. Black dots indicate individual sampling location. (b) Ringing and recapture locations of barn owls known to have flown out of (Emigrants) or into (Immigrants) Great Britain from 1910 to 2019, based on data courtesy of EURING. Lines simply connect two capture points and do not represent the actual path travelled by birds. Emigrant ringing locations in GB are coloured in blue, and recaptures in red. Immigrants into GB are coloured according to country of origin (orange – Belgium; green – Germany; blue – The Netherlands).

Analyses of capture-recapture data of ringed owls (N=80’083 individuals, from 1910 to 2019) revealed that all individuals ringed in Ireland (N=81 individuals) were recaptured in Ireland. As for GB, the vast majority (99.92%) of its ringed individuals (N=17’903) were also recaptured in GB and only 14 migrated out of the island: seven to Ireland (100% of this island’s immigrants) and seven to mainland Europe (Fig. 2b – Emigrants; Sup. Table 4a). In the opposite direction, GB received 21 individuals from the mainland (Fig. 2b - Immigrants), specifically from Belgium, the Netherlands and northern Germany (Sup. Table 4b). Of the immigrant birds, 19 were found dead, one severely injured with unknown fate, and one breeding. The latter was a female from the Netherlands, but the fate of its brood is not known. In the mainland, central European countries show considerably higher exchanges of individuals with each other (Sup. Table 4c) than with GB (Sup. Table 4b).

### Post-glacial species distribution

Habitat suitability projections for barn owls in the past showed that, at the time of the last glaciation, there was suitable land for barn owls outside of the known refugium of Iberia from a climatic perspective (Fig. 4c). Specifically, south of today’s British Isles there was a corridor of suitable land submerged nowadays, as well as along the south and western coasts of France, and a small cluster inland southern France. At the mid-Holocene (6’000 years BP), the coastline resembled that of present day, and the distribution of suitable habitat for barn owls resembled that of nowadays (Fig. 4c).

### Demographic inference

AIC and raw likelihood comparisons showed the Iberian origin B model to be the best at explaining the SFS of our dataset (Sup. Table 6; Fig. 4b). In this model, an ancestral insular lineage split from the mainland refugium lineage in Iberia fairly soon after the end of the glaciation, estimated at approximately 13’000 years ago (95% CI: 7’000-17’000 years BP; calculated with 3-year generation time). Only much later, the model predicted the split of the central EU population from PT at 4’000 years BP (95% CI: 1’000-5’000 years BP) and the separation between GB and IR at 1’200 years BP (95% CI: 220-2’200 years BP). Estimated effective population size was the largest in the PT population, followed by EU, GB and IR (Fig. 4b). Migration between populations was higher before these split than in recent times (Sup. Table 8; Ancestral vs Recent migration). Highest recent gene flow was observed from PT to EU, agreeing with Treemix’s first migration event (Sup. Fig. 5). Migration levels between the two islands and with the mainland were of a similar order of magnitude and less than half of that between mainland populations, consistent with the two barriers to gene flow identified by EEMS (Fig. 2a). Point estimates with 95% confidence intervals for all parameters of the best model are provided (Sup. Table 8), as well as single point estimates for the rest of the models (Sup. Table 7).

### Genome scans of colour-linked genes

Genome-wide scans revealed some high peaks of differentiation between populations, but none overlapped with the colour-linked candidate genes tested (Appendix 3). In particular, the *MC1R* region showed no particular sign of increased differentiation between pairs of populations, nor drop in diversity, with the exception of the known causal SNP between populations with different genotypes (Fig. 5b; Appendix 3).

## Discussion

Like most terrestrial species, barn owls are assumed to have colonized the British Isles after the last glaciation by crossing over Doggerland, a land bridge that connected GB to northern Europe. In continental Europe, barn owls display a marked latitudinal colour cline maintained through local adaptation (Antoniazza et al., 2010). However, in the British Isles they are conspicuously white in comparison to their nearest mainland counterparts questioning whether this is their source population. The currently held interpretation for their whiteness is a strong selection on this trait after colonisation. Here we provide evidence for a simpler explanation that does not require selection. Using whole-genome sequences and demographic simulations, we show that the colour disparity can be explained by the patterns in neutral genetic differentiation, resulting from an unexpected colonization route to the British Isles. We provide evidence for an early split of the insular lineage and low levels of gene flow with the mainland. Having found no evidence of selection on colour in the British Isles, it is plausible that this population has simply remained the white colour of its founders.

### Genetic isolation from the mainland

Our results based on whole genomes revealed genetic structure among western European barn owls despite shallow differentiation for a cosmopolitan bird (overall *F*_ST_ 0.035) and showed genome wide genetic isolation between the islands and the mainland, accompanied by low levels of gene flow and migration. On the mainland, Portugal displayed the highest levels of genetic diversity (Sup. Table 2) and the largest estimated population size (Figure 4b; Sup. Table 8), in accordance with its known role as a glacial refugium (Antoniazza et al., 2014). While forming its own population cluster (Figure 1b,c), we found evidence of considerable gene flow towards central Europe (Figure 2a, 4a,b; Sup. Table 8), consistent with a recent split between the two populations (less than 5’000 years BP; Figure 4a) and the relatively low differentiation between them (Sup. Table 3). This suggests that the Pyrenees are permeable to barn owl migration, unlike other higher and larger mountain ranges (Machado, Clément, Uva, Goudet, & Roulin, 2018). In central Europe, barn owl populations appear to be remarkably homogenous genetically, despite covering a large geographical and colour range (Figure 1, Sup. Table 3), in accordance with previous studies of continental Europe with traditional markers (Antoniazza et al., 2010), and supported by capture-recapture data that revealed high amounts of exchanges in central Europe (Sup. Table 4c).

Ireland and GB showed the lowest diversity and estimated effective population sizes in our study (Fig.4; Sup. Tables 2, 8). Barn owl populations of each island are genetically distinct from each other as well as from the mainland (Figure 1, 4a; Sup. Table 3). Genomic differentiation (Figure 1, 2a, 4a,b; Sup. Table 3) and capture-recapture data with only a handful of exchanges recorded in the last century (Figure 2b; Sup. Table 4a&b), suggest gene flow with the mainland is low. Specific analyses highlighted a barrier to gene flow extending from the Celtic Sea, through the English Channel to the North Sea (Figure 2a), effectively isolating the British Isles from the mainland.

Between the two islands, isolation appears to be recent (less than 2230 years BP; Figure 4a,b; Sup. Table 8), despite relatively high genetic differentiation (Sup. Table 3) likely exacerbated by an important effect of genetic drift in such small populations. There is little sign of current pervasive admixture in either direction (Figure 1c), consistent with the role of the Irish Sea as a strong barrier. However, there are records of owls from GB migrating into northern parts of Ireland (Figure 2b – Emigrants), the most easily accessible part of the island, while avoiding major water bodies by island-hopping from Scotland. Curiously, three of the individuals we sampled in Ireland for whole-genome sequencing (all sampled from found carcasses) appeared to be genetically from GB (Figure 1b,c), driving EEMS to place a gene flow barrier nearly along the political border between the two countries of Ireland instead of the sea (Figure 2a). Northern Ireland appears to be inhospitable for barn owls, at least in modern times, with only 1 to 3 pairs recorded per year in the whole country (*Barn Owl Report*, 2019). It could be acting as an extension of the sea barrier with the birds that fly in from GB being unable to find mates and thus not contributing to the genetic pool of the southern population, accentuating the differentiation between the two islands.

### Disparity in plumage colouration

Plumage colouration in barn owls, and the linked *MC1R* locus, follow a clinal distribution in continental Europe maintained by local adaptation (Antoniazza et al., 2010; Burri et al., 2016). Here, we formally establish that barn owls from the British Isles do not follow the continental latitudinal cline and are whiter than any continental population in Europe, including even Portugal (Figure 1a), confirming what was previously untested common knowledge among ornithologists. The rufous *MC1R* allele appears to be virtually absent in these populations in contrast to its close to 50% frequency at similar latitudes on the mainland, where dark morphs are positively selected (Figure 1a; Antoniazza et al. 2014; Burri et al. 2016). While genome-wide scans confirmed the important role of the known *MC1R* mutation in determining rufous colouration (Figure 5a), it appears to be restricted to the SNP variant itself and not the adjacent genomic regions (Figure 5b). Our results are consistent with previous studies that showed that carrying a single copy of the rufous allele is sufficient to ensure a darker phenotype, while individuals homozygous for the white allele can have a wide range of colouration (Burri et al., 2016; San-Jose et al., 2015).

This colour trait is likely polygenic, given that the known *MC1R* mutation explains only 30% of its variation (Burri et al., 2016; San-Jose et al., 2015) and its high heritability (Roulin & Dijkstra, 2003). Other loci could act in conjunction with a homozygous white *MC1R* to either produce whiter birds in GB or slightly darker morphs in Iberia. However, none of the other known colour-linked genes tested here explain how white owls homozygous for the white *MC1R* allele from Portugal reach darker phenotypes than those of the British Isles (Figure 1a, 5a; Appendix 3). Alternatively, it is conceivable that the phenotype we observe – colouration – simply reflects the pleiotropic effect of insular local adaptation on other linked cryptic traits. The melanocortin system regulates behaviour and physiology alongside the production of melanin, and associations between these traits are common among vertebrates (Ducrest, Keller, & Roulin, 2008; Roulin & Ducrest, 2011). Further work, potentially focusing on colour-varied populations to avoid the confounding factor of population structure could help elucidate the genetic basis of barn owl plumage colouration. If such other loci are found, it would be fascinating to investigate their distributions and interaction with *MC1R* along the continental colour cline as well as on the British Isles.

### Colonisation of the British Isles

Demographic simulations based on neutral sites showed that the British Isles were colonized from the glacial refugium in the Iberian Peninsula soon after the end of the glaciation (Figure 3b). This would have occurred while the British Isles were still connected to the mainland and the landmass extended considerably further south than today’s islands, following a corridor of suitable climatic conditions along the coast leading west (Figure 4c) completely separate from Doggerland. It is also possible that this corridor was already occupied by barn owls in a continuous population with Iberia before becoming isolated, as this species easily maintains high over-land gene flow (Figure 1b&c, 2a; Sup. Table 4c). Our wide confidence intervals make it hard to pin-point exactly the time of the actual split between the insular lineage from that of Iberia, but with the fast rise of sea levels and opening of the delta in the English Channel, the southern route to the islands would have been closed by 10’000 years BP (Lambeck, Rouby, Purcell, Sun, & Sambridge, 2014; Leorri, Cearreta, & Milne, 2012). Crucially, at this time prey would already be available in the form of voles, shrews, lemmings and bats (Montgomery et al., 2014). Once separated, the insular lineage underwent a long period of genetic drift, isolated from the mainland population in Iberia but homogenous within itself before splitting between the two islands (Figure 4a,b).

**Figure 3.**
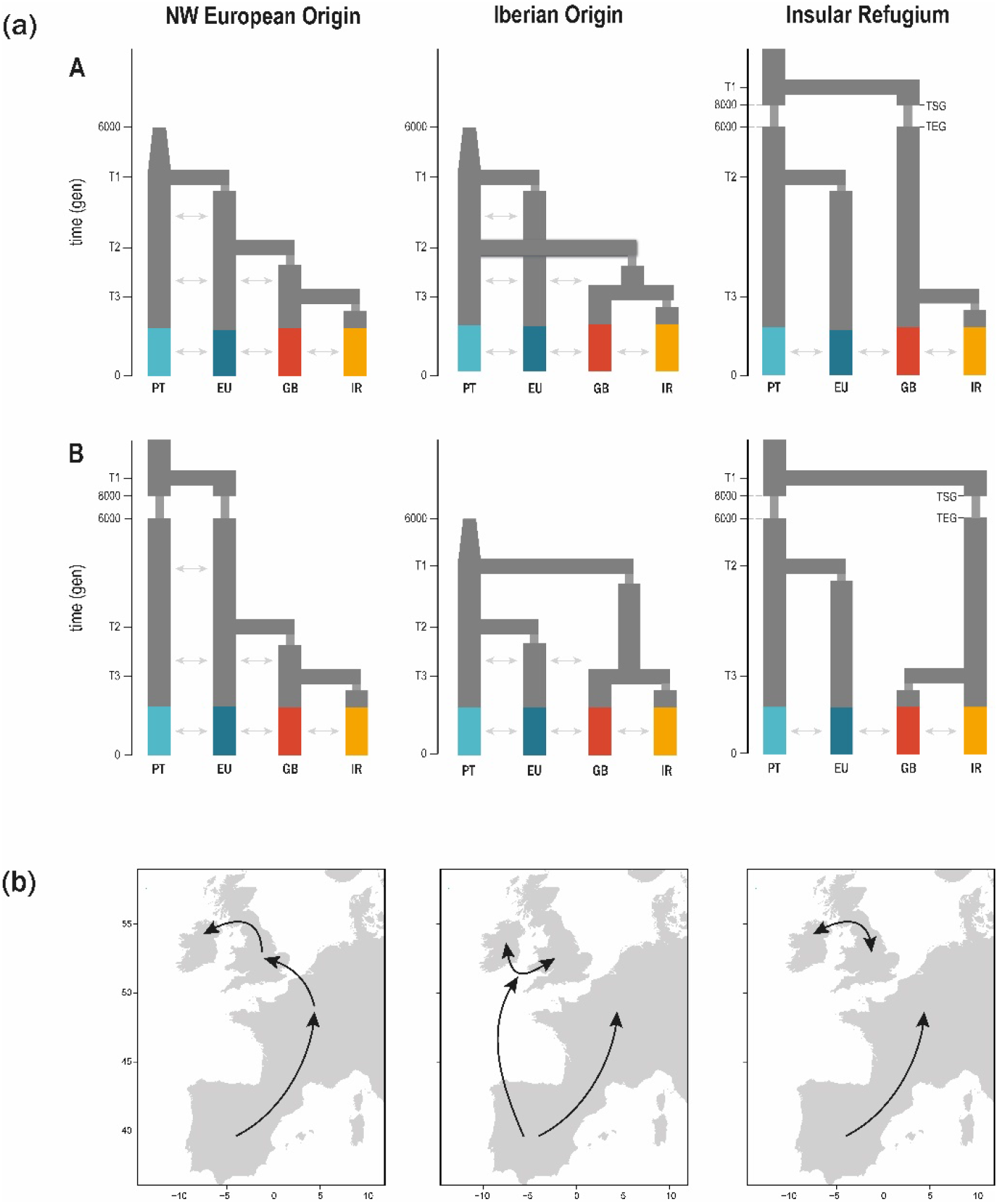
Hypothesized demographic scenarios for the colonization of the British Isles by barn owls. (a) Tested demographic scenarios for the colonization of the British Isles by barn owls. There are three main topologies – NW European Origin, Iberian Origin and Insular Refugium – each with two version (A & B; first and second line respectively). The four main genetic clusters in our dataset were used: Portugal (PT), Central Europe (EU), Great Britain (GB) and Ireland (IR). Population EU in this analysis is composed of individuals from FR and DK. Indicated times were fixed in the models (6’000 and 8’000 generations ago), and the remaining time parameters were inferred relative to them or to the event immediately before (e.g., T3 was bound between the present and T2). Cones depict post-glacial size increase and arrows gene flow between adjacent populations. In Insular Refugium topologies, TSG= time of start of glaciation in the insular lineage, TEG= time of end of glaciation in the insular lineage. (b) Schematic representation of the colonisation route to the British Isles for each scenario.

**Figure 4.**
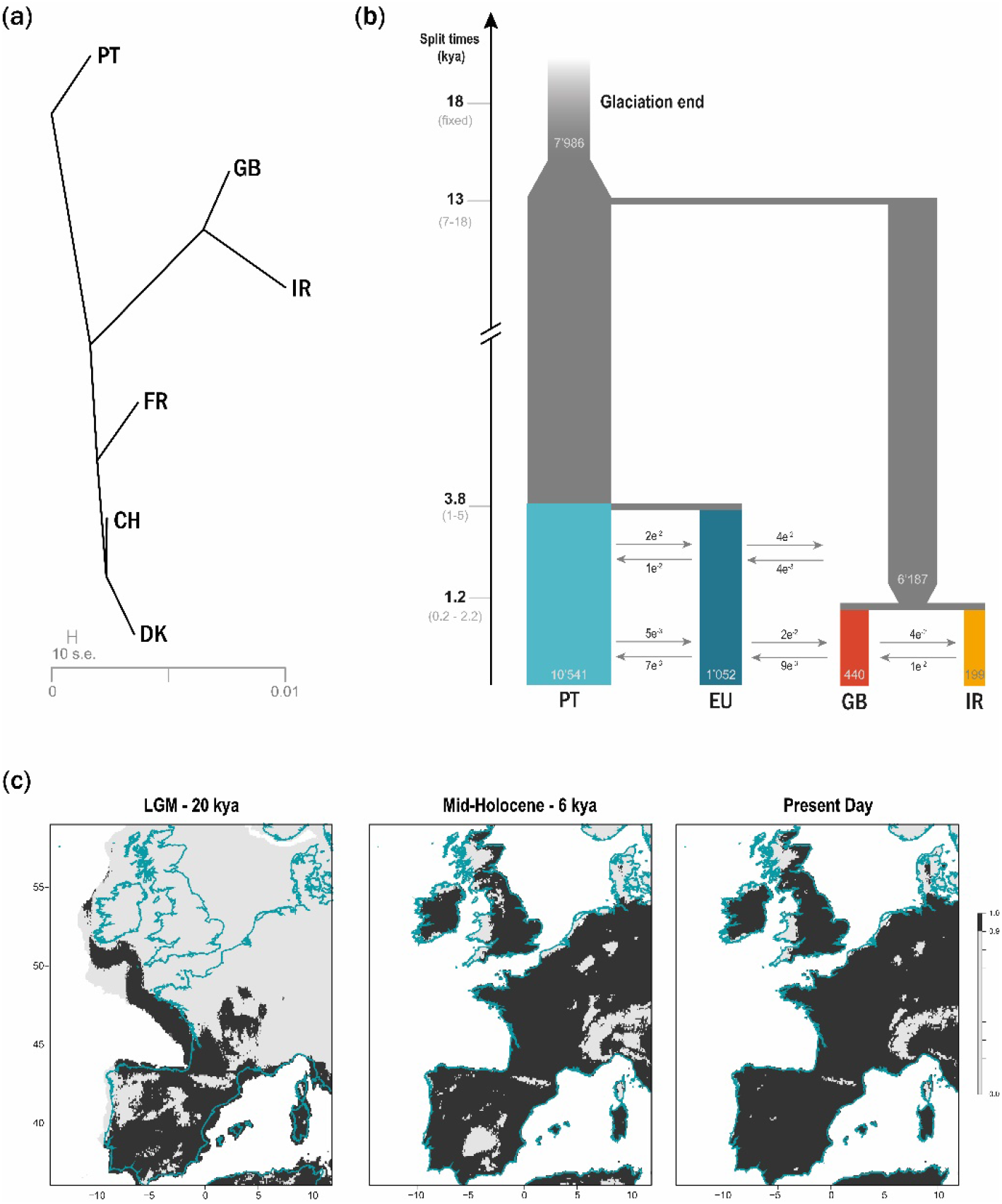
Demographic history of barn owls of the British Isles. (a) Treemix analysis with zero migration events. (b) Best supported demographic model for the colonisation of the British Isles as determined by fastsimcoal2. Time is indicated in thousands of years, with a 3-year generation time, confidence intervals at 95% are given between brackets. Population sizes (haploid) are shown inside each population bar; arrows indicate forward-in-time migration rate and direction. Population EU in this analysis is composed of individuals from FR and DK. (c) Species distribution model of barn owls projected into past conditions – last glacial maximum (20’000 years BP) and mid-Holocene (6’000 years BP) – compared to today’s distribution. Only locations with high suitability in at least 90% model averaging are coloured in dark grey. Below that threshold cells were considered as unsuitable (lightest grey shade on the graph). Modern coastline is shown in blue.

On the mainland, central European populations split genetically from the Iberian refugium much later (less than 5’000 years BP). Large population sizes and high overland gene flow (Figure 4b; Sup. Table 8) might thus have maintained low differentiation for a long period of time, but also climatic conditions north of the Pyrenees may have taken longer to become favourable. The latter hypothesis would further counter the traditional point of view of Doggerland as the point of arrival for barn owls, as they could have not yet reached such high latitudes before Doggerland submerged 8’000 years BP. Intriguingly, our demographic model predicts high migration from GB into central Europe between the splits of the latter with Iberia and between the two islands (Figure 4b), which appears unlikely with all land bridges submerged at this point (less than 5’000 years BP). It is possible that the migration rate was inflated as the model did not allow for gene flow between the ancestral insular and mainland populations before the first split and thus forced all migration to occur in a short time interval (Figure 3).

In light of the inferred demographic history, barn owls of the British Isles would have inherited their whiteness from their source mainland population, the refugium in the Iberian Peninsula, and kept it through small population size, genetic drift and low gene flow. Although it is conceivable that some copies of the rufous *MC1R* allele were present in the founding insular population, similar to its frequency in Iberia (1%; Figure 1a), in the absence of strong positive selection in the insular environment, it could have disappeared through genetic drift given the small effective sizes (Figure 4b; Sup. Table 8). Thus, the selective pressure that renders the rufous colour and allele adaptive in northern continental Europe (Antoniazza et al., 2010; Burri et al., 2016), may be absent in the British Isles. Still, we cannot rule out that gene flow with the mainland is too weak and over too short a period of time to offer selection sufficient variation in the British Isles to increase the frequency of imported rufous alleles. If, conversely, the white morph was positively selected on the islands – potentially explaining its purer shade – we would have expected to find extended haplotypic differentiation when comparing it to the white mainland birds, which we did not (Figure 5; Appendix 3). Therefore, it appears the white insular morph can be most parsimoniously explained by relaxation or absence of selective pressure in contrast to the mainland. Such a pattern is actually common among insular birds which, due to relaxed selection, tend to display less colourful plumage than their mainland counterparts (Doutrelant et al., 2016; Grant, 1965), as also observed in the barn owl worldwide (Romano, Séchaud, & Roulin, 2021).

**Figure 5.**
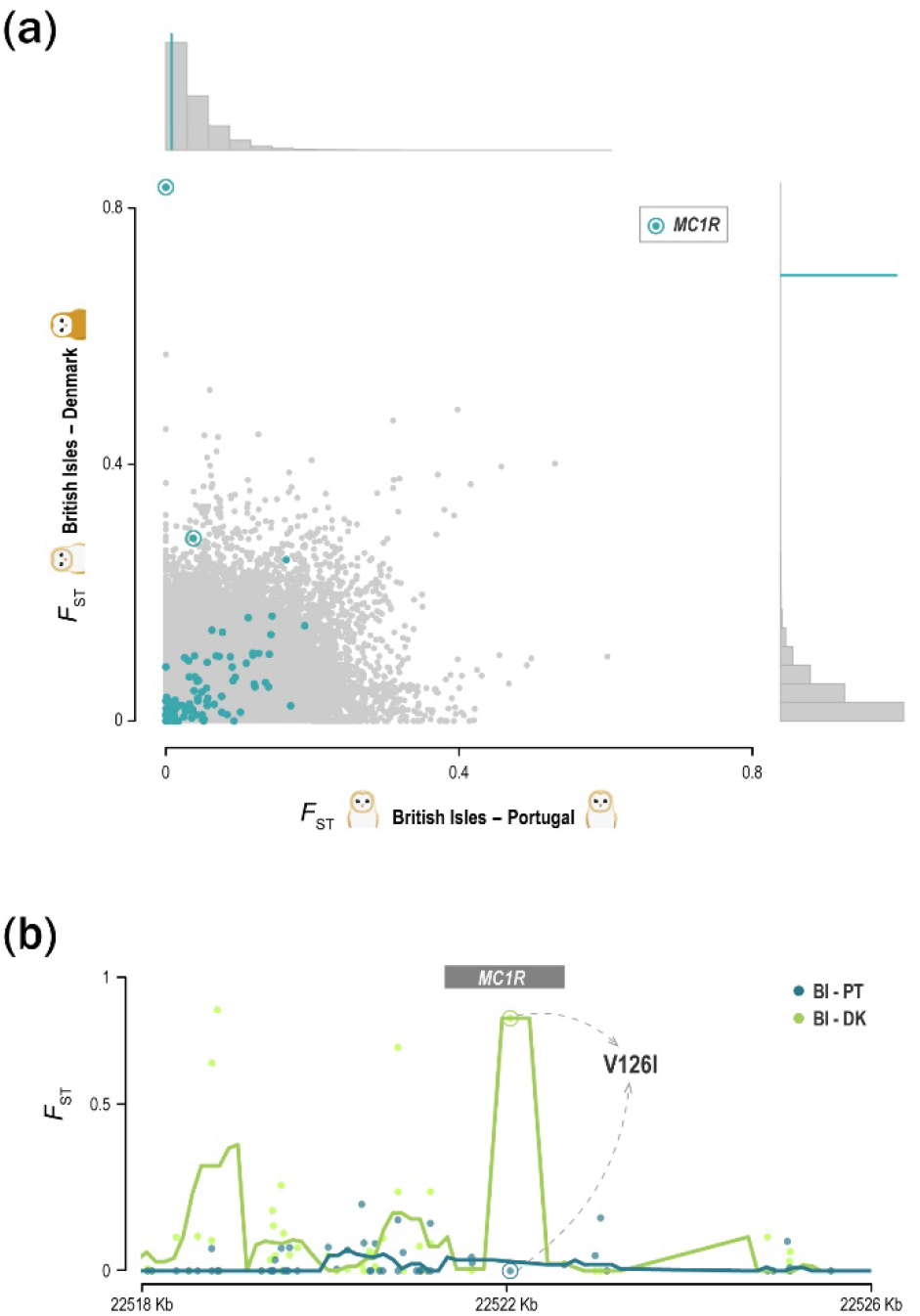
Differentiation at the colour-linked locus V126I of the *MC1R* gene between differently coloured barn owl populations in Europe. (a) Genome-wide *F*_ST_ values per window (in grey, 20Kbp windows with 5Kbp steps), between two white barn owl populations on the horizontal axis – British Isles (BI) and Portugal (PT) – and between one white and one rufous on the vertical axis – BI and Denmark (DK). The distribution of each axis is shown on the histograms. Blue dots indicate the *F*_ST_ at windows containing the tested colour-linked genes. Windows containing the *MC1R* are encircled, and their mean is shown with the blue line on the histograms. (b) *F*_ST_ per site (dots) around the *MC1R* gene (grey box). Lines show the mean over sliding windows (500bp with 100bp step), for the same comparisons as above: BI and PT in blue; BI and DK in green. Circled dots indicate the V126I locus in both comparisons.

This early history of colonisation of the British Isles inferred here from whole-genome sequences and supported by SDM projections on past climatic features is apparently unique among terrestrial vertebrates, but it is far from the first to deviate from the most common colonisation route over Doggerland (e.g. Boston et al., 2015; Kelly et al., 2010; Snell et al., 2005; Stewart & Lister, 2001; Teacher et al., 2009) or to indicate an earlier colonisation than generally assumed (Martínková et al., 2007; McDevitt et al., 2020). The case of the stoat (*Mustela erminea*) is particularly interesting as it was found to have had an isolated glacial refugium also in now submerged land southwest of today’s French coastline on the Bay of Biscay (Figure 4c – LGM; Martínková et al. 2007). From there they reached Ireland very early as the temperatures started rising but, as the Celtic Sea opened 15’000 years BP, only colonized GB much later over Doggerland (Martínková et al., 2007). The key difference between the two cases lies in the fact that barn owls maintained a homogenous population between GB and Ireland through flight.

### Conclusion

Our study demonstrates that barn owls followed a highly uncommon post-glacial colonisation route to the British Isles. Likely taking advantage of the since submerged suitable habitat on the Bay of Biscay, barn owls reached the islands much earlier than expected from this southern point. The inferred demographic history could explain the whiteness of these populations through a combination of founder effect and low gene flow, and without the need to invoke selective pressures. We contend high quality population genomic data associated with species distribution hindcasting will reveal an unusual demographic history and post-glacial colonization for many non-model species. We wonder how often an intuitive selective explanation for a conspicuous phenotype could turn out to be the result of purely neutral processes.

## Acknowledgements

We are grateful to the following institutions and individuals that provided samples or aided in sampling to our study: Barn Owl Trust (Devon, UK), BirdWatch Ireland (Kilcoole, IE), National University of Ireland (Galway, IE), The European Barn Owl Network, CHENE Association (Seine-Maritime, FR), Burke Museum of Natural History and Culture (Washington, USA), San Diego Natural history Museum (California, USA), the late Richard F. Shore, Emy Guibault, Sylvain Antoniazza and Reto Burri. We thank Céline Simon, Guillaume Dumont, Luis San-José, Valérie Ducret and Kathleen Salin for their valuable assistance with molecular work. Finally, we thank Laurent Excoffier and Nina Marchi for their assistance with demographic inference in Fastsimcoal2, Olivier Broennimann for his advice and review of the SDM, and John Pannell for insightful review of the manuscript. This study was funded by the Swiss National Science Foundation with grants 31003A_138180 & 31003A_179358 to JG and 31003A_173178 to AR.

## Data Accessibility

The new refence genome for European barn owl (*Tyto alba*) has been deposited at DDBJ/ENA/GenBank under the accession JAEUGV000000000, and the corresponding PacBio reads in the BioProject PRJNA694553. The raw Illumina reads for the whole-genome sequenced individuals are available in BioProject PRJNA700797. Colour and MC1R data are provided in Appendix 1.

## Author Contributions

APM, TC, AR, JG designed this study; APM produced whole-genome resequencing libraries; APM, TC conducted the analyses; ALD, MD produced the new reference genome; CI, EB, NG assembled it; TC identified coding regions; KD, RL, JL, HDM, LP and DR provided samples to the study; APM led the writing of the manuscript with input from all authors.

