## Supporting Information for "Unexpected post-glacial colonisation route explains the white colour of barn owls (*Tyto alba*) from the British Isles"

### Contents

Supporting Methods ... 2

Whole-genome resequencing ... 2

Data preparation and SNP calling ... 2

Post-glacial species distribution ... 3

Maximum-likelihood demographic inference with *fastsimcoal2* ... 5

Supporting Methods References ... 7

Supporting Tables ... 9

Supporting table 1 ... 9

Supporting table 2 ... 9

Supporting table 3 ... 9

Supporting table 4 ... 10

Supporting table 5 ... 11

Supporting table 6 ... 11

Supporting table 7 ... 12

Supporting table 8 ... 13

Supporting Figures ... 14

Supporting Figure 1 ... 14

Supporting Figure 2 ... 15

Supporting Figure 3 ... 16

Supporting Figure 4 ... 16

Supporting Figure 5 ... 17

### Supporting Methods

#### Whole-genome resequencing

The whole genomes of 63 individuals (61 European and 2 outgroups) were sequenced in this study. Their sex was determined with molecular markers (Py, Ducrest, Duvoisin, Fumagalli, & Roulin, 2006) prior to sequencing. DNA quality and fragmentation were assessed with Fragment Analyzer™ (Advanced Analytical Technologies Inc.). Sample Purification Beads (SPB; Illumina, California, USA) were used to remove fragments smaller than 500 bp prior to library preparation of samples that showed high spread of DNA fragment size. Further, the intensity of the initial mechanical fragmentation step (Covaris, Woburn, MA, USA) was adjusted based on Fragment Analyzer profiles to promote homogenous library sizes. Individually tagged 100bp TruSeq DNA PCR-free libraries (Illumina) were prepared according to manufacturer's instructions. Whole-genome resequencing was performed on multiplexed libraries with Illumina HiSeq 2500 high-throughput paired-end sequencing technology at the Lausanne Genomic Technologies Facility (GTF, University of Lausanne, Switzerland).

#### Data preparation and SNP calling

Raw reads were trimmed with Trimommatic v.0.36 (Bolger, Lohse, & Usadel, 2014) for Illumina adapters, and minimum sequence length of 70 bp. BWA-MEM v.0.7.15 (H. Li & Durbin, 2009) was used to map the trimmed reads to the newly generated reference barn owl genome. Despite our libraries being PCR-free, potentially duplicate reads were marked with Picard-tools v2.9.0 (http://broadinstitute.github.io/picard) MarkDuplicates per run and per library.

Base quality score recalibration (BQSR) was performed following the iterative approach recommended for non-model species for which a set of "true variants" is not available, using high-confidence calls on the un-calibrated calls in a bootstrap-fashion to achieve convergence of the quality of variant calls. Here, a first calling of high-confidence variants was done with a combination of GATK's HaplotypeCaller and GenotypeGVCF v.4.1.3 and ANGSD v.0.921

(Korneliussen, Albrechtsen, & Nielsen, 2014). Both sets of calls were filtered for a maximum of 5% missing data, individual depth (DP > 10 and DP < 30), mapping quality (MQ > 40) and minor allelic frequency (MAF > 0.02). The intersect of the two call sets was used as the known set of variants to run BQSR a first time with BaseRecalibrator and ApplyBQSR in GATK v.4.1.3. The calibrated output was used to recall variants with GATK and ANGSD as above, and the intersect was used on a second round of recalibration. The results were similar to the previous run suggesting convergence had been achieved as observed in other bird genomics studies (Burri et al., 2015). Thus, the recalibrated calls obtained in the first round were kept for the remainder of the pipeline. Following BQSR, sequence variants were called with GATK's HaplotypeCaller and GenotypeGVCFs v.4.1.3 from the recalibrated bam files.

Genotype calls were filtered for downstream analyses using a hard-filtering approach as proposed for non-model organisms, using GATK and VCFtools (Danecek et al., 2011). Calls were removed if they presented: low individual quality per depth (QD < 5), extreme coverage (800 > DP > 1800), mapping quality (MQ < 40 and MQ > 70), extreme hetero or homozygosity (ExcessHet > 20 and InbreedingCoeff > 0.9) and high read strand bias (FS > 60 and SOR > 3). We filtered further at the level of individual genotype by removing calls for which up to 5% of genotypes had low quality (GQ < 20) and extreme coverage (GenDP < 10 and GenDP > 40). Lastly, we kept only bi-allelic sites with less than 5% of missing calls across individuals yielding a dataset of 6'721'999 SNP. For analyses of neutral population structure and demography, an exact Hardy-Weinberg test was used to remove sites that significantly departed ($p<0.05$) from the expected equilibrium using the R (R Development Core Team, 2016) package HardyWeinberg (Graffelman, 2015; Graffelman & Morales-Camarena, 2008).

#### Post-glacial species distribution

We built species distribution models (SDM) using Maximum Entropy Modelling (MaxEnt), a presence-only based modelling tool, to identify the regions of high habitat suitability for barn owls at the last glacial maximum (LGM, 20’000 years BP). Current climatic variables for the Western Palearctic (Sup. Fig. 2) were extracted from the WorldClim database (Hijmans, Cameron, Parra, Jones, & Jarvis, 2005) at 5 arc min resolution using the R package rbioclim (Exposito-Alonso, 2017). The chosen set of variables represents the climatic conditions experienced by the species through the year and were filtered to be correlated at less than 0.8. Retained variables were: mean diurnal range (bio2), minimum temperature of coldest month (bio6), temperature annual range (bio7), mean temperature of wettest quarter (bio8), precipitation seasonality (bio15), precipitation of driest quarter (bio17) and precipitation of coldest quarter (bio19).

To determine which combination of feature and regularization multiplier use in the model to optimise the prediction without over complexifying the model, we built models with linear, quadratic and hinge features, and models with a range (1 to 5) of regularization multipliers. The best combination of feature and regularization multiplier based on the corrected AIC (as recommended by Warren & Seifert, 2011) was achieved with a quadratic model with 1 as regularization multiplier (Sup. Table 8). We ran 100 independent maxent models, omitting 25% of the data during training to test the model. To avoid geographic bias due to different sampling effort in the distribution area of the species, we randomly extracted 1000 presence points within the IUCN distribution map (BirdLife International, 2019) for each model run (Fourcade, 2016).

Predictive performances of the models were evaluated on the basis of the area under the curve of the receiver operator plot (AUC) of the test data. For all models with an AUC higher than 0.8 (considered a good model; Swets 1988; Li et al. 2020), the output of maxent was transformed into a binary map of suitability by assuming that a cell was suitable when its mean suitability value was higher than the mean value of the 10% test presence threshold. This conservative threshold allows to omit all regions with habitat suitability lower than the suitability values for the lowest 10% of occurrence records. We averaged the values of the models for each cell, and only cells suitable in 90% of the models are presented as such in the map.

All models were then projected to the mid-Holocene (6'000 years BP) and LGM (20'000 years BP) conditions extracted from WorldClim at the same resolution as current data. When projecting to past climates, the multivariate environmental similarity surface (MESS) approach (Elith, Kearney, & Phillips, 2010) was used to assess whether models were projected into climatic conditions different from those found in the calibration data. Cells with climatic conditions outside the distribution used to build the model were considered as unsuitable for barn owls (0 attributed to cell with negative MESS) as we intended to highlight only the highly suitable regions. For each timepoint, the results of the models were merged and transformed into a binary map as for the current data.

To determine which combination of feature and regularization multiplier use in the model to optimise the prediction without over complexifying the model, we built models with linear, quadratic and hinge features, and models with a range (1 to 5) of regularization multipliers. The best combination of feature and regularization multiplier based on the corrected AIC (as recommended by Warren & Seifert, 2011) was achieved with a quadratic model with 1 as regularization multiplier (Sup. Table 5).

When projecting to past climates, the multivariate environmental similarity surface (MESS) approach (Elith et al., 2010) was used to assess whether models were projected into climatic conditions different from those found in the calibration data. Cells with climatic conditions outside the distribution used to build the model were considered as unsuitable for barn owls (0 attributed to cell with negative MESS) as we intended to highlight only the highly suitable regions.

### Maximum-likelihood demographic inference with *fastsimcoal2*

#### *Data preparation*

To build a set of variants approaching neutrality, we kept only autosomal SNPs found outside of genic regions and CpG mutations (Pouyet, Aeschbacher, Thiéry, & Excoffier, 2018). The sites were filtered to include no missing data and to within two thirds of the standard deviation of the mean coverage to ensure homogeneity. To determine the ancestral state of the SNP using a parsimony approach, the genomes of the two outgroups were used as outgroups based on the *Tytonidae* phylogenetic tree (Uva, Päckert, Cibois, Fumagalli, & Roulin, 2018). Sites for which it was impossible to attribute a state based on the available outgroups were discarded (868 sites).

*Demographic inference*

For each run of each of the six tested demographic scenarios, the following options were set: -n 500000 (number of coalescent simulations), -M and -L 50 (estimate the parameters from the SFS with 50 expectation-maximization (EM) cycles to estimate parameters). As we do not currently have a good estimation of the barn owl mutation rate, the end of the glaciation (rounded to 6000 generations ago, 18'000 years BP with a 3-year generation time) was fixed and all other parameters were scaled relative to it using the -0 option (based solely on polymorphic sites).

In the non-parametric bootstrapping of the best-fitting model, a block-bootstrap approach was employed as suggested by the authors to account for LD (Excoffier, Dupanloup, Huerta-Sánchez, Sousa, & Foll, 2013; Excoffier & Foll, 2011). As such, the SNPs were divided into 100 blocks of similar size and then 100 bootstrap datasets were generated by sampling with replacement 100 blocks each, so as to obtain the same number of SNPs as the real dataset. For each bootstrapped SFS, 50 independent parameter inferences were run under the best-fitting scenario out of the six tested. Due to computational constraints, bootstrap runs were performed with only 10 EM cycles, an approach that has been previously used and described as conservative (Malaspinas et al., 2016). The highest maximum-likelihood run of each scenario was used to estimate 95% CI of all parameters.

### Supporting Methods References


BirdLife International, . (2019). The IUCN Red List of Threatened Species. Version 6.2. Retrieved November 19, 2020, from BirdLife International website: https://dx.doi.org/10.2305/IUCN.UK.2019-3.RLTS.T22688504A155542941.en

Bolger, A. M., Lohse, M., & Usadel, B. (2014). Trimmomatic: A flexible trimmer for Illumina sequence data. *Bioinformatics*, *30*(15), 2114–2120. doi: 10.1093/bioinformatics/btu170

Burri, R., Nater, A., Kawakami, T., Mugal, C. F., Olason, P. I., Smeds, L., ... Ellegren, H. (2015). Linked selection and recombination rate variation drive the evolution of the genomic landscape of differentiation across the speciation continuum of *Ficedula* flycatchers. *Genome Research*, *25*, 1656–1665. doi: 10.1101/gr.196485.115

Danecek, P., Auton, A., Abecasis, G., Albers, C. A., Banks, E., DePristo, M. A., ... Durbin, R. (2011). The variant call format and VCFtools. *Bioinformatics*, *27*(15), 2156–2158. doi: 10.1093/bioinformatics/btr330

Elith, J., Kearney, M., & Phillips, S. (2010). The art of modelling range-shifting species. *Methods in Ecology and Evolution*, *1*(4), 330–342. doi: 10.1111/j.2041-210x.2010.00036.x

Excoffier, L., Dupanloup, I., Huerta-Sánchez, E., Sousa, V. C., & Foll, M. (2013). Robust Demographic Inference from Genomic and SNP Data. *PLoS Genetics*, *9*(10), 1003905. doi: 10.1371/journal.pgen.1003905

Excoffier, L., & Foll, M. (2011). fastsimcoal: A continuous-time coalescent simulator of genomic diversity under arbitrarily complex evolutionary scenarios. *Bioinformatics*, *27*(9), 1332–1334. doi: 10.1093/bioinformatics/btr124

Exposito-Alonso, M. (2017). *rbioclim: Improved getData function from the raster R package to interact with past, present and future climate data from worldclim.org*. Retrieved from github.com/MoisesExpositoAlonso/rbioclim

Fourcade, Y. (2016). Comparing species distributions modelled from occurrence data and from expert-based range maps. Implication for predicting range shifts with climate change. *Ecological Informatics*, *36*, 8–14. doi: 10.1016/j.ecoinf.2016.09.002

Graffelman, J. (2015). Exploring Diallelic Genetic Markers: The {HardyWeinberg} Package. *Journal of Statistical Software*, *64*(3), 1–23. Retrieved from http://www.jstatsoft.org/v64/i03/

Graffelman, J., & Morales-Camarena, J. (2008). Graphical tests for Hardy-Weinberg Equilibrium based on the ternary plot. *Human Heredity*, *65*(2), 77–84.

Hijmans, R. J., Cameron, S. E., Parra, J. L., Jones, P. G., & Jarvis, A. (2005). Very high resolution interpolated climate surfaces for global land areas. *International Journal of Climatology*, *25*(15), 1965–1978. doi: 10.1002/joc.1276

Korneliussen, T. S., Albrechtsen, A., & Nielsen, R. (2014). ANGSD: Analysis of Next Generation Sequencing Data. *BMC Bioinformatics*, *15*(1), 1–13. doi: 10.1186/s12859-014-0356-4

Li, H., & Durbin, R. (2009). Fast and accurate short read alignment with Burrows-Wheeler transform. *Bioinformatics*, *25*(14), 1754–1760. doi: 10.1093/bioinformatics/btp324

Li, Y., Li, M., Li, C., & Liu, Z. (2020). Optimized maxent model predictions of climate change impacts on the suitable distribution of cunninghamia lanceolata in China. *Forests*, *11*(3), 302. doi: 10.3390/f11030302

Malaspinas, A.-S., Westaway, M. C., Muller, C., Sousa, V. C., Lao, O., Alves, I., … Willerslev, E. (2016). A genomic history of Aboriginal Australia. *Nature*, *538*(7624), 207–214. doi: 10.1038/nature18299

Pouyet, F., Aeschbacher, S., Thiéry, A., & Excoffier, L. (2018). Background selection and biased gene conversion affect more than 95% of the human genome and bias demographic inferences. *ELife*, *7*. doi: 10.7554/eLife.36317

Py, I., Ducrest, A.-L., Duvoisin, N., Fumagalli, L., & Roulin, A. (2006). Ultraviolet reflectance in a melanin-based plumage trait is heritable. *Evolutionary Ecology Research*, *8*(3), 483–491.

R Development Core Team. (2016). R: A language and environment for statistical computing. *R Foundation for Statistical Computing, Vienna, Austria*. R Foundation for Statistical Computing, Vienna, Austria. Retrieved from https://www.r-project.org/

Swets, J. A. (1988). Measuring the accuracy of diagnostic systems. *Science*, *240*(4857), 1285–1293. doi: 10.1126/science.3287615

Uva, V., Päckert, M., Cibois, A., Fumagalli, L., & Roulin, A. (2018). Comprehensive molecular phylogeny of barn owls and relatives (Family: Tytonidae), and their six major Pleistocene radiations. *Molecular Phylogenetics and Evolution*, *125*, 127–137. doi: 10.1016/j.ympev.2018.03.013

Warren, D. L., & Seifert, S. N. (2011). Ecological niche modeling in Maxent: The importance of model complexity and the performance of model selection criteria. *Ecological Applications*, *21*(2), 335–342. doi: 10.1890/10-1171.1

### Supporting Tables

**Supporting table 1** – Summary of the sampling scheme for the different analyses in this study. Full sample detail available in Appendix 1.

| Population | Abbrev. | N Colour | N MC1R | N WGS | N FSC |
|---|---|---|---|---|---|
| Ireland | IR | 44 | 44 | 12 | 8 |
| Great Britain | GB | 67 | 68 | 15 | 8 |
| France | FR | 48 | 48 | 5 | 5 |
| Denmark | DK | 88 | 88 | 10 | 3 |
| Switzerland | CH | 65 | 65 | 10 | 0 |
| Portugal | PT | 70 | 71 | 9 | 8 |
| Total | | 382 | 384 | 61 | 32 |

Number of samples for: N Colour – colour comparison analysis; N MC1R – MC1R genotyping; N WGS – whole genome re-sequencing and population genomics analyses; N FSC – demographic inference with *fastsimcoal2*.

**Supporting table 2** – Lineage genetic diversity for of 61 European barn owls. Individuals were grouped into the four main lineages identified in sNMF (Fig. 1 & Sup. Fig. 3).

| Lineage | Abbrev. | N | Priv. Alleles | $H_O$ (SD) | $F_{IS}$ (SD) |
|---|---|---|---|---|---|
| Ireland | IR | 9 | 84061 | 0.157 (0.004) | -0.023 (0.027) |
| Great Britain | GB | 17 | 93144 | 0.158 (0.011) | 0.029 (0.068) |
| Central Europe | EU | 25 | 191666 | 0.174 (0.010) | -0.004 (0.057) |
| Portugal | PT | 9 | 784099 | 0.188 (0.008) | -0.012 (0.041) |

N – number of individuals; Priv. Alleles – private alleles accounting for different population sizes; $H_O$ – observed heterozygosity and its standard deviation; $F_{IS}$ – inbreeding coefficient.

**Supporting table 3** – Pairwise Weir & Cockerham's $F_{ST}$ between barn owl populations in Western Europe. Above the diagonal, a heat map provides a visual representation of the $F_{ST}$ values given below the diagonal.

| | IR | GB | FR | DK | CH | PT |
|---|---|---|---|---|---|---|
| IR | | | | | | |
| GB | 0.037 | | | | | |
| FR | 0.064 | 0.041 | | | | |
| DK | 0.061 | 0.038 | 0.016 | | | |
| CH | 0.054 | 0.032 | 0.012 | 0.005 | | |
| PT | 0.060 | 0.041 | 0.021 | 0.022 | 0.017 | |

**Supporting table 4** – Summary of barn owl capture-recapture events in Great Britain (GB) and Central Europe from 1910 to 2019, courtesy of EURING. Table shows number and percentages (%) of **(a)** emigrant and **(b)** immigrant owls to GB. Table **(c)** shows exchanges of individuals between countries in Central Europe; number of owls indicates migrants from the country on the row towards the country in column; background heatmap represents the values on top; two last columns give the number of ringed birds per country and how many of those emigrated elsewhere. BL= Belgium; CH=Switzerland; DE=Germany; DK=Denmark; FR=France; NL=Netherlands.

(a)

| Country | Total immigrants to this country | Immigrants from GB | % immigrants from GB |
|---|---|---|---|
| Belgium | 338 | 1 | 0.30 |
| France | 1506 | 1 | 0.07 |
| Germany | 1364 | 1 | 0.07 |
| Netherlands | 1107 | 1 | 0.09 |
| Spain | 39 | 3 | 7.69 |
| Ireland | 4 | 4 | 100 |
| Northern Ireland | 4 | 3 | 75 |

(b)

| Country | Total ringed owls | Total emigrant owls | % emigrant owls | Emigrant owls to GB | % emigrant owls to GB |
|---|---|---|---|---|---|
| Belgium | 6166 | 1070 | 0.28 | 3 | 0.049 |
| Germany | 20816 | 1493 | 0.35 | 5 | 0.024 |
| Netherlands | 25849 | 1177 | 1.19 | 13 | 0.050 |

(c)

| → | BL | CH | DE | DK | FR | NL | N ringed | Emigrants |
|---|---|---|---|---|---|---|---|---|
| BL | | 3 | 80 | 3 | 342 | 616 | 7786 | 1118 |
| CH | 3 | | 502 | 0 | 518 | 6 | 7862 | 1074 |
| DE | 61 | 66 | | 136 | 486 | 424 | 24245 | 1522 |
| DK | 1 | 0 | 44 | | 0 | 4 | 1003 | 57 |
| FR | 9 | 6 | 36 | 0 | | 8 | 1318 | 67 |
| NL | 251 | 6 | 652 | 10 | 104 | | 29487 | 1177 |

**Supporting table 5**– Comparison of SDM model fit. AICc is reported for the multiple combinations of feature (linear, quadratic, hinge) and Beta multiplier (1 to 5).

| | 1 | 2 | 3 | 4 | 5 |
|---|---|---|---|---|---|
| **Linear** | 23953.4 | 23994.8 | 24037.1 | 24086.8 | 24125.8 |
| **Quadratic** | 23950.8 | 24002.6 | 24042.7 | 24095.2 | 24148.5 |
| **Hinge** | 24099.1 | 24185.4 | 24241.2 | 24304.4 | 24340.2 |

**Supporting table 6** – Likelihood and AIC of the demographic models tested with *fastsimcoal2*. Three main model topologies were tested, each with two versions (Figure 3). Models are sorted from best to worst according to the estimated likelihoods.

| Model Type | Model | Est. Lhood | Δ Lhood | AIC | Δ AIC |
|---|---|---|---|---|---|
| Colour-based | B | -6410181 | 4342 | 29520023 | 0 |
| Refugia | A | -6410377 | 4537 | 29520926 | 904 |
| Colour-based | A | -6410439 | 4599 | 29521216 | 1193 |
| Refugia | B | -6410674 | 4834 | 29522291 | 2269 |
| Stepping-Stone | B | -6410811 | 4971 | 29522935 | 2912 |
| Stepping-Stone | A | -6410865 | 5025 | 29523172 | 3150 |

Est. Lhood – Maximum-likelihood estimated for the simulated SFS per demographic model; Δ Lhood – difference between the likelihood of the simulated and observed SFS; Δ AIC – delta AIC

**Supporting table 7** – Parameter ranges and point estimate inferred for the demographic model tested with *fastsimcoal2*. Model "Iberian origin B" – identified as the best fitting model – is given in Sup. Table 8. All range distributions were uniform except for bottlenecks (†) which were log-uniform. When a parameter was absent in a model, the case was left blank.

| | | NW European | | Iberian | Insular Refugium | |
|---|---|---|---|---|---|---|
| **Parameter** | **Ranges** | **A** | **B** | **A** | **A** | **B** |
| *Current Population Sizes (haploid)* | | | | | | |
| PT | 10000 – 4e5 | 58059 | 14700 | 10518 | 35058 | 61402 |
| EU | 1000 - 3.5e5 | 1018 | 4097 | 1011 | 1007 | 5314 |
| GB | 100 - 25000 | 1008 | 139 | 137 | 200 | 1712 |
| IR | 10 - 2500 | 421 | 221 | 232 | 179 | 390 |
| *Ancestral Population Sizes (haploid)* | | | | | | |
| PT in glac | 200 – 2e5 | 39179 | | 6637 | | |
| Ancestral island | 200 – 2e5 | | | 11277 | | |
| PT before glac | 1000 - 1e6 | | 16066 | | 8603 | 13846 |
| GB before glac | 1000 - 50000 | | | | 41926 | 23204 |
| EU before glac | 1000 – 3.5e5 | | 207200 | | | |
| *Times of Divergence (generations)* | | | | | | |
| T1 | | 4875 | 12758 | 3656 | 11705 | 30398 |
| T2 | | 3635 | 256 | 3055 | 2843 | 2469 |
| T3 | | 304 | 204 | 288 | 1219 | 3318 |
| TSG | 8000 - 9000 | | | | 8452 | 8148 |
| TEG | 3000 - 6000 | | | | 3409 | 4482 |
| *Current Migration (flow is backwards in time)* | | | | | | |
| EU → PT | 0 - 0.05 | 0.004 | 0.0001 | 0.005 | 0.007 | 0.001 |
| PT → EU | 0 - 0.05 | 0.011 | 0.005 | 0.011 | 0.004 | 0.001 |
| GB → EU | 0 - 0.05 | 0.004 | 0.058 | 0.054 | 0.039 | 0.005 |
| EU → GB | 0 - 0.05 | 0.011 | 0.002 | 0.013 | 0.005 | 0.0007 |
| IR → GB | 0 - 0.05 | 0.014 | 0.032 | 0.027 | 0.042 | 0.018 |
| GB → IR | 0 - 0.05 | 0.007 | 0.042 | 0.042 | 0.025 | 0.0001 |
| *Older Migration (2nd level from present in Sup. Fig. 1)* | | | | | | |
| EU → PT | 0 - 0.05 | 0.002 | 0.018 | 0.0004 | | |
| PT → EU | 0 - 0.05 | 0.035 | 0.014 | 0.001 | | |
| GB → EU | 0 - 0.05 | 0.006 | 0.032 | 0.002 | | |
| EU → GB | 0 - 0.05 | 0.021 | 0.049 | 0.030 | | |
| *Ancestral Migration (oldest)* | | | | | | |
| EU → PT | 0 - 0.05 | 0.002 | 0.004 | 0.008 | | |
| PT → EU | 0 - 0.05 | 0.00002 | 0.022 | 0.011 | | |
| *Instbot- bottleneck intensity at diverge †* | | | | | | |
| T1 | 0.01 - 0.5 | 0.052 | 0.006 | 0.006 | 0.009 | 0.003 |
| T2 | 0.01 - 0.5 | 0.004 | 0.139 | 0.696 | 0.037 | 0.019 |
| T3 | 0.01 - 0.5 | 0.010 | 0.016 | | 0.044 | 0.110 |
| *Glacation Bottleneck size (haploid) †* | | | | | | |
| PT | 0.01 - 0.5 | | 295 | | 2747 | 23223 |
| EU | 0.01 - 0.5 | | 490 | | | |
| GB | 0.01 - 0.5 | | | | 8 | 39 |

**Supporting table 8** – Parameter point estimates and 95% confidence interval for the best demographic model – Iberian origin B. Parameter names correspond to Figure 3. Times of divergence are in years calculated with a generation time of 3 years. Migration rates and number of individuals are given forward in time.

| Parameter | Point Estimate | Lower Limit CI | Upper Limit CI |
|---|---|---|---|
| *Current Population Sizes (haploid)* | | | |
| PT | 10541 | 10501 | 364413 |
| EU | 1052 | 1005 | 2382 |
| GB | 440 | 131 | 1838 |
| IR | 199 | 138 | 1071 |
| *Ancestral Population Sizes (haploid)* | | | |
| Iberia glacial period | 7986 | 5451 | 137520 |
| Ancestral pop to GB & IR | 6187 | 2112 | 20742 |
| *Times of Divergence (years)* | | | |
| Split GB - PT | 12940 | 7316 | 17812 |
| Split EU - PT | 3822 | 945 | 4984 |
| Split IR - GB | 1149 | 224 | 2229 |
| *Current Migration Rate (2Nm)* | | | |
| PT → EU | 0.005 (52) | 0.0009 (10) | 0.0089 (93) |
| EU → PT | 0.0074 (8) | 0.0038 (4) | 0.0158 (17) |
| EU → GB | 0.0152 (16) | 0.0003 (0.3) | 0.0548 (58) |
| GB → EU | 0.0093 (4) | 0.0051 (2) | 0.0206 (9) |
| GB → IR | 0.0361 (16) | 0.0019 (1) | 0.0473 (21) |
| IR → GB | 0.0103 (2) | 0.0011 (0.2) | 0.0438 (9) |
| *Ancestral Migration Rate (2Nm)* | | | |
| PT → EU | 0.0029 (262) | 0.0002 (2) | 0.0524 (552) |
| EU → PT | 0.0248 (3) | 0.0031 (3) | 0.0585 (62) |
| EU → GB | 0.0001 (0.1) | 0.0004 (0.4) | 0.0159 (17) |
| GB → EU | 0.03 (186) | 0.0121 (75) | 0.0633 (392) |
| *Instant Bottlenecks (N)* | | | |
| Split GB - PT | 0.015 (65) | 0.288 (4) | 0.01 (98) |
| Split EU - PT | 0.018 (56) | 0.032 (31) | 0.002 (535) |

### Supporting Figures

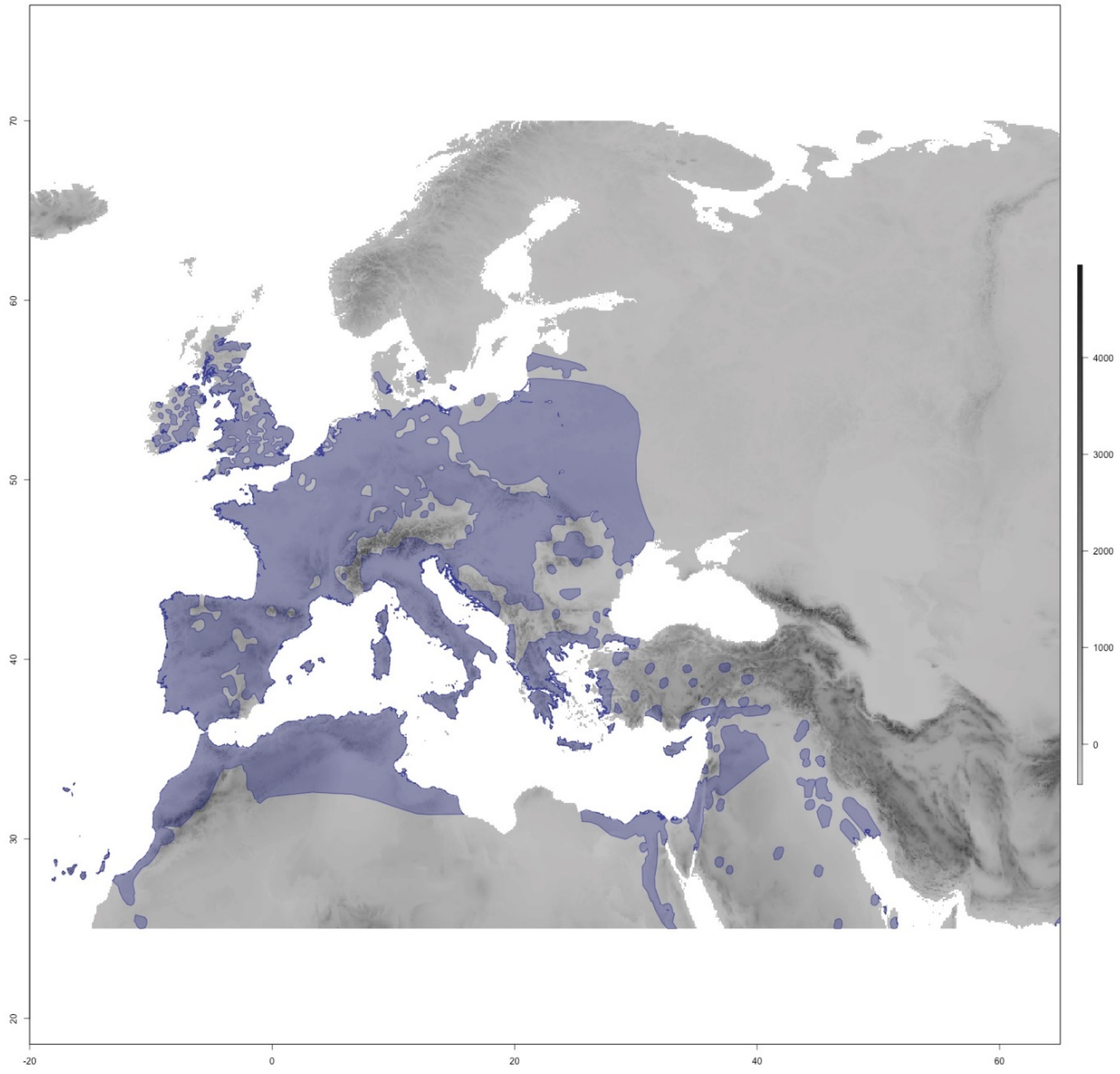


**Supporting Figure 1** – The map in grey represents the area considered for producing the Species Distribution Model (SDM) for the barn owl; shading denotes altitude according to the scale. The current distribution of barn owls is plotted atop the map in purple (data from IUCN: BirdLife International 2019). Random presence points were extracted within this distribution for the SDM.

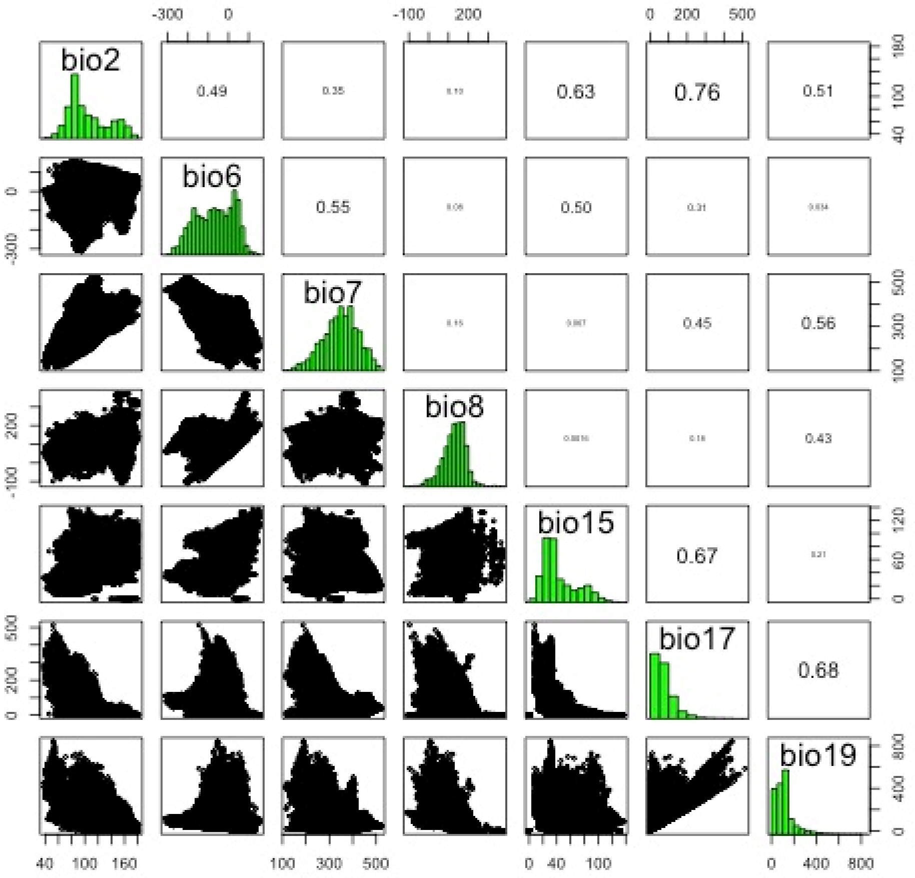


**Supporting Figure 2** – Pairwise correlation between retained climatic variables to produce the SDM. Only variables correlated at less than 0.8 were kept in the models, namely: Mean Diurnal Range (Bio2), Min Temperature of Coldest Month (Bio6), Temperature Annual Range (Bio7), Mean Temperature of Wettest Quarter (Bio8), Precipitation Seasonality (Bio15), Precipitation of Driest Quarter (Bio17) and Precipitation of Coldest Quarter (Bio19).

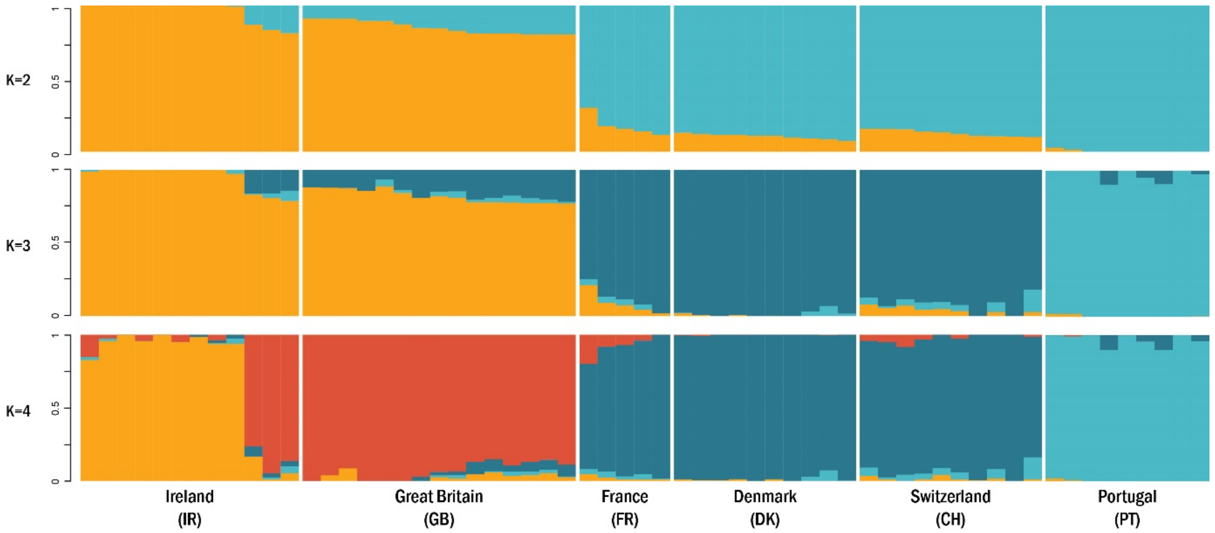


**Supporting Figure 3** – Individual clustering estimated by sNMF for K 2 to 4 lineages. Each vertical bar represents one individual, and the colours represent the relative contributions of each genetic lineage K.

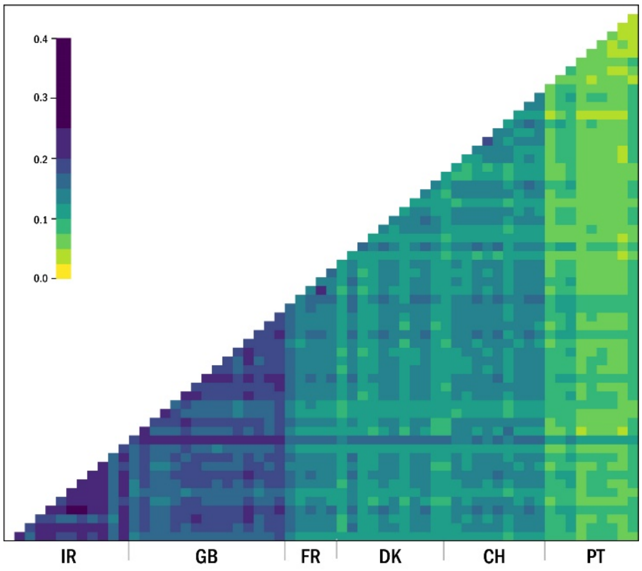


**Supporting Figure 4** – Individual relatedness (β) matrix. Grey lines separate the populations. Population abbreviations follow Sup. Fig. 3.

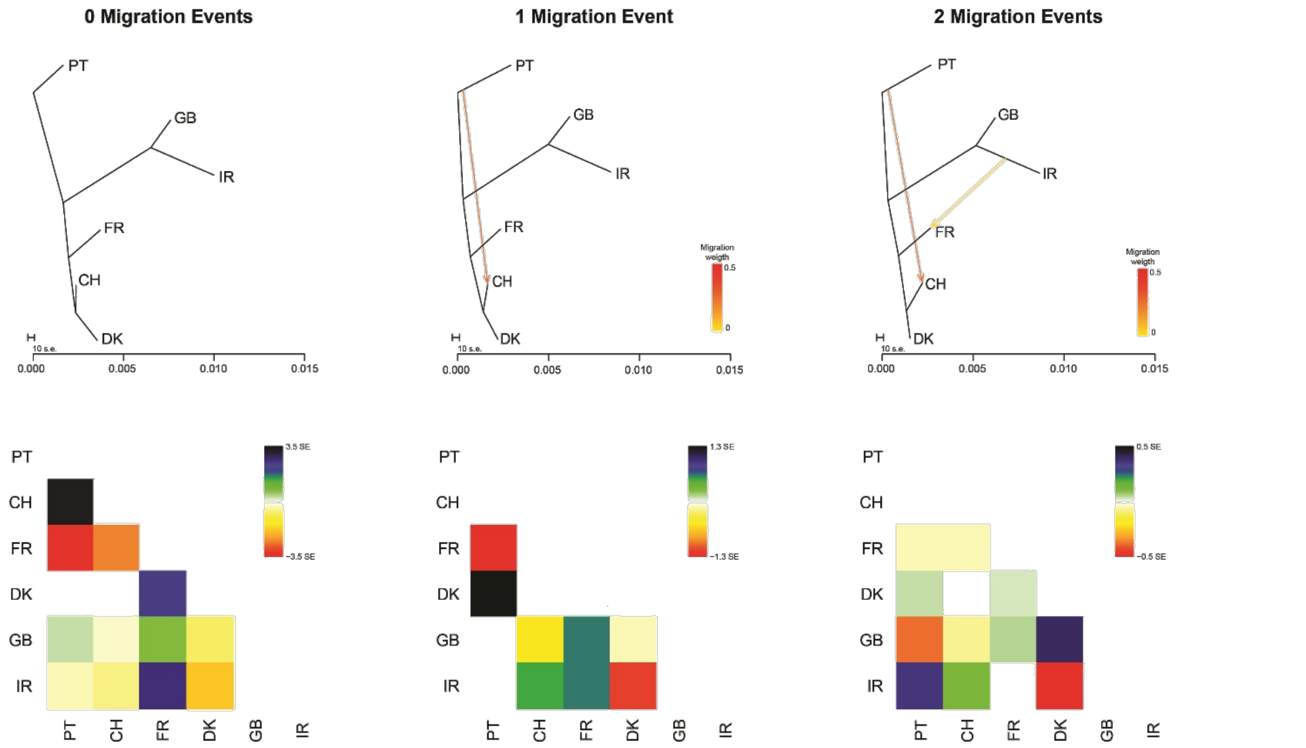


**Supporting Figure 5** – Treemix results for 0 to 2 migration events. Highest likelihood runs are depicted, with the corresponding matrix of standard errors. With two migration events there was no topology convergence between replicates; the topology of the best run shown here was only present in 4 out of 10 runs.
