## Appendix2 for "Unexpected post-glacial colonisation route explains the white colour of barn owls (*Tyto alba*) from the British Isles"

### APPENDIX 2 – New reference genome of European barn owl (*Tyto alba*)

*Extraction of high molecular weight genomic DNA*

A fresh blood sample was collected from a young Swiss male barn owl (M040663). High molecular weight (HMW) genomic DNA was extracted using an agarose plug method as described by Zhang et al. (2012). In brief, red cells were counted. For 10 µg of DNA-agarose plug of 100 µl, 10 µl of red cells washed in PBS and resuspended in 190 µl of PBS were added to 200 µl of 1% (wt/vol) low melt agarose (Low gelling temperature agarose, Life Technologies) at 45°C. 100 ul of the mix was applied to the disposable plug molds (Bio-Rad). The plugs were transferred to falcon tubes and cells digested with 0.5M EDTA pH 9.3, 1 % (wt/vol) sodium lauryl sarcosine and 0.3 mg/ml of proteinase K (Promega) for 24h at 50°C with slow shaking 40 rpm. The plugs were then washed once in 50 mM EDTA pH 8.0, then 3x in 10 mM Tris-HCl pH 8.0, 1 mM EDTA (TE 1x) with 0.1 mM PMSF, 3x TE 1x buffer, each wash for 1 hour on ice. Then following the Bionano protocol (bionano Genomics), each plug was transferred to 1.5 ml eppendorf tube, excess of liquid was wiped out, the plug was quickly spin down and place at 65°C for 10 min, then transferred to 42°C for 5 min. One ul of 1 U/ul of beta-agarase (Bioconcept) was added and the tube incubated for 45 min at 42°C. The DNA was then dialysed on a nitrocellulose 0.1 um millipore membrane (Merck) for 45 min in TE 1x buffer. Viscous HMW DNA was recovered in a 1.5 ml eppendorf tube and kept at 4°C. After 24h, the DNA was homogenized by pipetting up down with wide-bore tips and quantified by Qubit. A femto pulsefield was conducted to control for DNA small fragments. On a pulse field gel DNA appears above 2.2 Mbp to 225 kbp.

*Optical mapping library preparation*

In silico digestion with the previous genome (Ducrest et al., 2020) was used to find the best enzyme combination for the optical mapping. Direct labeling with the non-nicking enzyme DLE-1,

(5’-CTTAAG-3’recognition site) and the nicking enzyme Nt.BspQ1 (5’-GCTCCTTN1/N4-3’) produced 17.6 and 15.1 labels per 100 kbp of genomic DNA. 750 ng of genomic DNA were subjected to direct labeling and 300 ng of DNA to nick-label-repair-stain reactions following exactly the Bionano protocols (bionano Genomics, San Diego, USA) at the Functional Genomics Center of Zurich (University of Zurich, Switzerland). At the end the labeled DNA was quantified with Qubit and loaded into a nanochannel array of a Saphyr Chip (bionano Genomics) and run by electrophoresis each into a compartment of the Saphyr system.

*Pacbio library preparation and Sequencing*

High molecular weight DNA was sheared with Megaruptor (Diagenode, Denville, NJ, USA) to obtain 80kb fragments. After shearing the DNA size distribution was checked on a Fragment Analyzer (Advanced Analytical Technologies, Ames, IA, USA). A SMRTbell library was prepared with five µg of the DNA with the PacBio SMRTbell Express Template Prep Kit 2.0 (Pacific Biosciences, Menlo Park, CA, USA) according to the manufacturer's recommendations. The resulting library was size selected on a BluePippin system (Sage Science, Inc. Beverly, MA, USA) for molecules larger than 35 kb that were sequenced on 12 SMRT cells 1M with v3.0/v3.0 chemistry and diffusion loading on a PacBio Sequel platform (Pacific Biosciences, Menlo Park, CA, USA) at 600 min movie length.

*Assembly*

Sequencing produced 7.3 million long reads with a total sum length of unique single molecules of 135 Gbp (N50 > 31Kb), an approximative 108X coverage for a 1.25Gb genome (135/1.25 = 103.84). Reads for the 12 SMRT cells have been deposited at DDBJ/ENA/GenBank under BioProject PRJNA694553. Reads were assembled using pb-assembly workflow from the PacBio Assembly Tool Suite (https://github.com/PacificBiosciences/pb-assembly) with default parameters. FALCON (Chin et al., 2016) was used to produce the primary assembly that yielded

10'814 contiguous primary contigs, a mixed of phased haplotypes and collapsed haplotype regions. Haplotype reconstruction was performed using FALCON-Unzip v.3 (Chin et al., 2016) resulting in 478 unzipped primary contigs partially phased, and 1736 fully phased haplotigs which represented divergent haplotypes.

The resulting contigs were assembled into scaffolds using Bionano Solve v.3.4.1 (Bionano Genomics, USA) with two optical mapping runs: DLE1 with nickase recognition site *cttaag* and BspQ1 with nickase recognition site *gctcttc*. Each Bionano run was assembled into contigs using the manufacturer's default pipeline and using the unzip contigs as hint for the rough assembly autonoise step of the pipeline (-r and -R parameters of pipelineCL.py). The 2 runs were then combined with the unzip contigs to produce the finished scaffolded assembly using Bionano's two-enzyme hybrid scaffold pipeline (runTGH.R) following the manufacturer's instructions. The resulting assembly was composed of 70 scaffolds, considerably closer to the barn owl’s karyotype of 46 chromosomes than the previously available references. See Appendix 2 Table 1 for the full assembly metrics.

**Appendix 2 Table 1** – Assembly metrics of the new barn owl reference genome, and comparison to the previously available genome (Ducrest et al. 2020).

| Parameter | New Genome | Ducrest 2020 |
|---|---:|---:|
| Length (bp) | 1’249’867’532 | 1’219’191’878 |
| Nb scaffolds | 70 | 21’509 |
| Longest scaffold (bp) | 91’687’297 | 22’155’979 |
| Shortest scaffold (bp) | 43’623 | 500 |
| N50 (bp) | 36’032’128 | 4’615’526 |
| L50 | 13 | 72 |
| N90 (bp) | 15’349’174 | 556’444 |
| L90 | 33 | 350 |
| N’s (bp) | 51’362’034 | 9’580’001 |

**Appendix 2 Table 3** – BUSCO scores of the assembly

| Category | N | % |
|---|---|---|
| Complete BUSCOs (C) | 8079 | 96.9 |
| *Complete and single-copy BUSCOs (S)* | 8056 | 96.6 |
| *Complete and duplicated BUSCOs (D)* | 23 | 0.3 |
| Fragmented BUSCOs (F) | 67 | 0.8 |
| Missing BUSCOs (M) | 192 | 2.3 |
| Total BUSCO groups searched | 8338 | 100 |

*Identification of repeated sequences and coding regions*

RepeatModeler v.1.0.11 (Smit & Hubley, 2008-2015) and RepeatMasker v.4.0.7 (Smit, Hubley, & Green, 2013-2015) were used to assess repeats and low complexity regions of the genome with default parameters. RepeatModeler, a combination of three de-novo repeat finding programs (RECON, RepeatScout and LtrHarvest/Ltr_retriever) that produce a high-quality library of transposable elements families, identified 122 families of repeated elements. Based on this library, RepeatMasker was used to screen the reference genome, identifying 7.26% of the assembly as interspersed repeats and 1.53% as low complexity DNA sequences (Appendix 2 Table 2).

**Appendix 2 Table 2** – Repetitive elements identified by RepeatMasker in the new reference genome.

| Type of repetitive element | Nb of elements | Length (bp) | % of sequence |
|---|---|---|---|
| SINEs | 2745 | 401711 | 0.03 |
| *ALUs* | 0 | 0 | 0 |
| *MIRs* | 0 | 0 | 0 |
| LINEs | 72891 | 30575904 | 2.45 |
| *LINE1* | 0 | 0 | 0 |
| *LINE2* | 0 | 0 | 0 |
| *L3/CR1* | 72891 | 30575904 | 2.45 |
| LTR elements | 4518 | 5587338 | 0.45 |
| *ERVL* | 1575 | 2102299 | 0.17 |
| *ERVL-MaLRs* | 0 | 0 | 0 |
| *ERV_classI* | 1702 | 1520708 | 0.12 |
| *ERV_classII* | 777 | 882367 | 0.07 |
| DNA elements | 0 | 0 | 0 |
| *hAT-Charlie* | 0 | 0 | 0 |
| *TcMar-Tigger* | 0 | 0 | 0 |
| Unclassified | 81555 | 35130478 | 2.81 |
| Total interspersed repeats | | 71695431 | 5.74 |
| Small RNA | 0 | 0 | 0 |
| Satellites | 1 | 224 | 0 |
| Simple repeats | 377585 | 15568066 | 1.25 |
| Low complexity | 65926 | 3543479 | 0.28 |

Red v.05.22.2015 (Girgis, 2015) was used to mark all repetitive regions in the genome through machine learning, soft masking 36.6% of the assembly. Six previously published mRNAseq libraries (Ducrest et al., 2020) were download from the European Bioinformatics Institute European Nucleotide Archive (accession number ERP115928, from doi.org/10.1002/ece3.5991). Reads were trimmed for adapter using Trimommatic v.0.36 (Bolger, Lohse, & Usadel, 2014), and then mapped to the masked reference using tophat2 v.2.1.1 (Kim et al., 2013) both with default settings. To identify coding regions, we applied the Braker2 pipeline v.2.0.1 (Barnett, Garrison, Quinlan, Strömberg, & Marth, 2011; Brůna, Hoff, Lomsadze, Stanke, & Borodovsky, 2020; Buchfink, Xie, & Huson, 2015; Gotoh, 2008; Hoff,

Lange, Lomsadze, Borodovsky, & Stanke, 2016; Hoff, Lomsadze, Borodovsky, & Stanke, 2019; Iwata & Gotoh, 2012; Li et al., 2009; Lomsadze, Burns, & Borodovsky, 2014; Lomsadze, Ter-Hovhannisyan, Chernoff, & Borodovsky, 2005; Stanke, Diekhans, Baertsch, & Haussler, 2008; Stanke, Schöffmann, Morgenstern, & Waack, 2006) on the soft masked version of the reference genome produced by Red. Braker2 is a combination of GeneMark-EP+ and AUGUSTUS, trained from RNA-Seq and/or protein homology information. We fed Braker2 with RNA-seq and protein data from OrthoDB v.101 restricted to proteins from Aves family (taxid 8782), yielding 19829 coding regions (Appendix 2 Table 3).

**Appendix 2 Table 3** – BUSCO scores of identified coding regions.

| Category | N | % |
|---|---|---|
| Complete BUSCOs (C) | 7046 | 84.5 |
| *Complete and single-copy BUSCOs (S)* | 7013 | 84.1 |
| *Complete and duplicated BUSCOs (D)* | 33 | 0.4 |
| Fragmented BUSCOs (F) | 677 | 8.1 |
| Missing BUSCOs (M) | 615 | 7.4 |
| Total BUSCO groups searched | 8338 | 100 |
