## Appendix3 for "Unexpected post-glacial colonisation route explains the white colour of barn owls (*Tyto alba*) from the British Isles"

### APPENDIX 3 – Genome-wide scans and focus on colour-linked genes

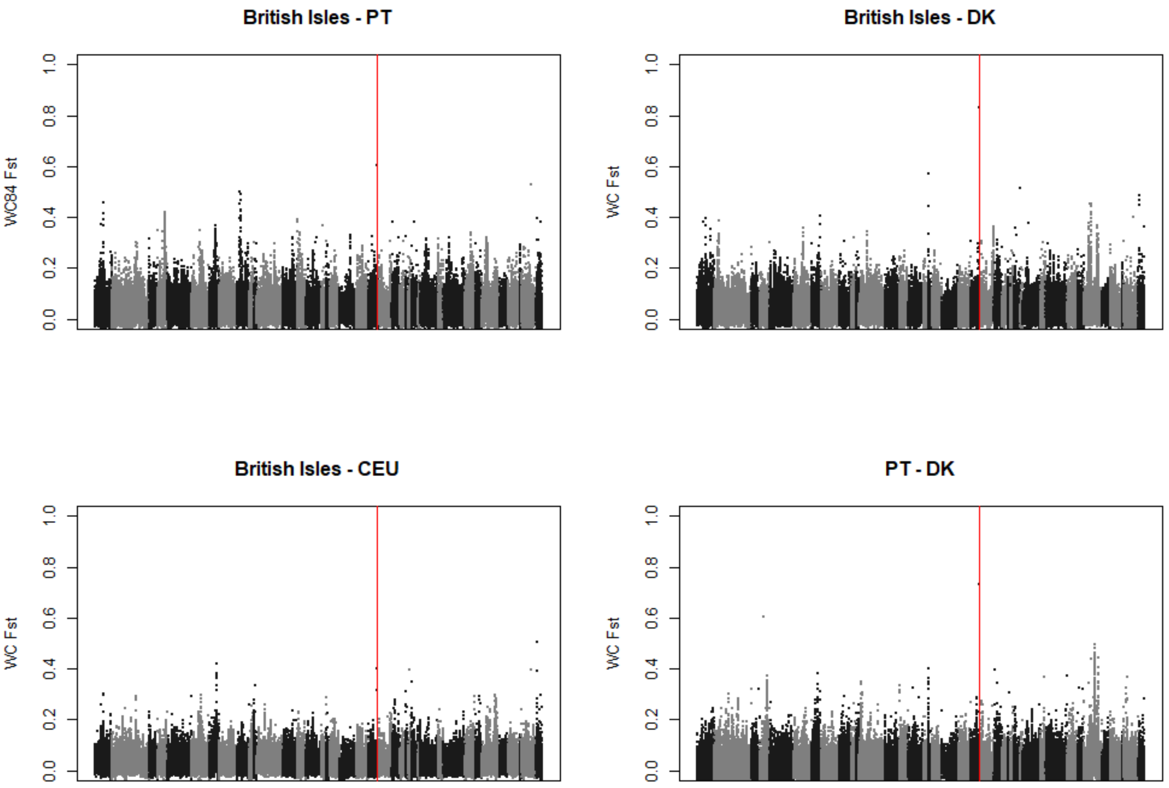


**Appendix 3 Figure 1** - Genome-wide $F_{ST}$ scans. between differently coloured populations, averaged over 20 kbp sliding windows in 5 kbp step. Alternation in colours indicates change of scaffold. The red line shows the position of *MC1R* in the genome.

In addition to *MC1R*, we mapped 22 other autosomal genes (Appendix 3 Table 1) shown to be linked to melanin colouration in other organisms onto the barn owl genome. Our goal was to see if, in the absence of any signal at *MC1R*, any of these genes showed signals of particular differentiation between plumage colour morphs. The figures of the following pages are organised as follows: each page concerns one gene (or specific exons of a gene) indicated on the title of the left-hand side plots. Plots on the left show pairwise $F_{ST}$ between pairs of populations indicated on the title, and on the right nucleotide diversity (π), both estimated over 5kbp sliding windows in

1kbp step. Brit – British Isles, includes Great Britian and Ireland; PT – Portugal; DK – Denmark; CEU – Central Europe, includes France and Switzerland.

**Appendix 3 Table 1** – List of 23 candidate genes analysed in this study. The references indicated are examples of each gene's link to melanin-based colouration.

| # | Gene | Reference |
|---|---|---|
| 1 | AGRP | (Li et al. 2011) |
| 2 | ASIP-ACD | (San-Jose et al. 2017) |
| 3 | ATP6V1C2 | (Schweizer et al. 2016) |
| 4 | CORIN | (Bourgeois et al. 2016) |
| 5 | CREB1 | (San-Jose et al. 2017) |
| 6 | CTHRC1 | (Zheng et al. 2020) |
| 7 | DCT | (San-Jose et al. 2017) |
| 8 | EDNRB2 | (Li et al. 2015) |
| 9 | GPER1 | (Natale et al. 2016) |
| 10 | GPR143 | (Shi et al. 2012) |
| 11 | GR | (Béziers et al. 2019) |
| 12 | KIT | (San-Jose et al. 2017) |
| 13 | MC1R | (San-Jose et al. 2017) |
| 14 | MFSD12 | (Aguillon et al. 2021) |
| 15 | MGRN1 | (Walker and Gunn 2010) |
| 16 | MITF-M | (Li et al. 2012) |
| 17 | MR | (Béziers et al. 2019) |
| 18 | OCA2 | (Kratochwil et al. 2019) |
| 19 | PCSK2 | (San-Jose et al. 2017) |
| 20 | POMC | (Li et al. 2011) |
| 21 | RAB38 | (Abolins-Abols et al. 2018) |
| 22 | SOX10 | (Li et al. 2011) |
| 23 | TYR | (San-Jose et al. 2017) |

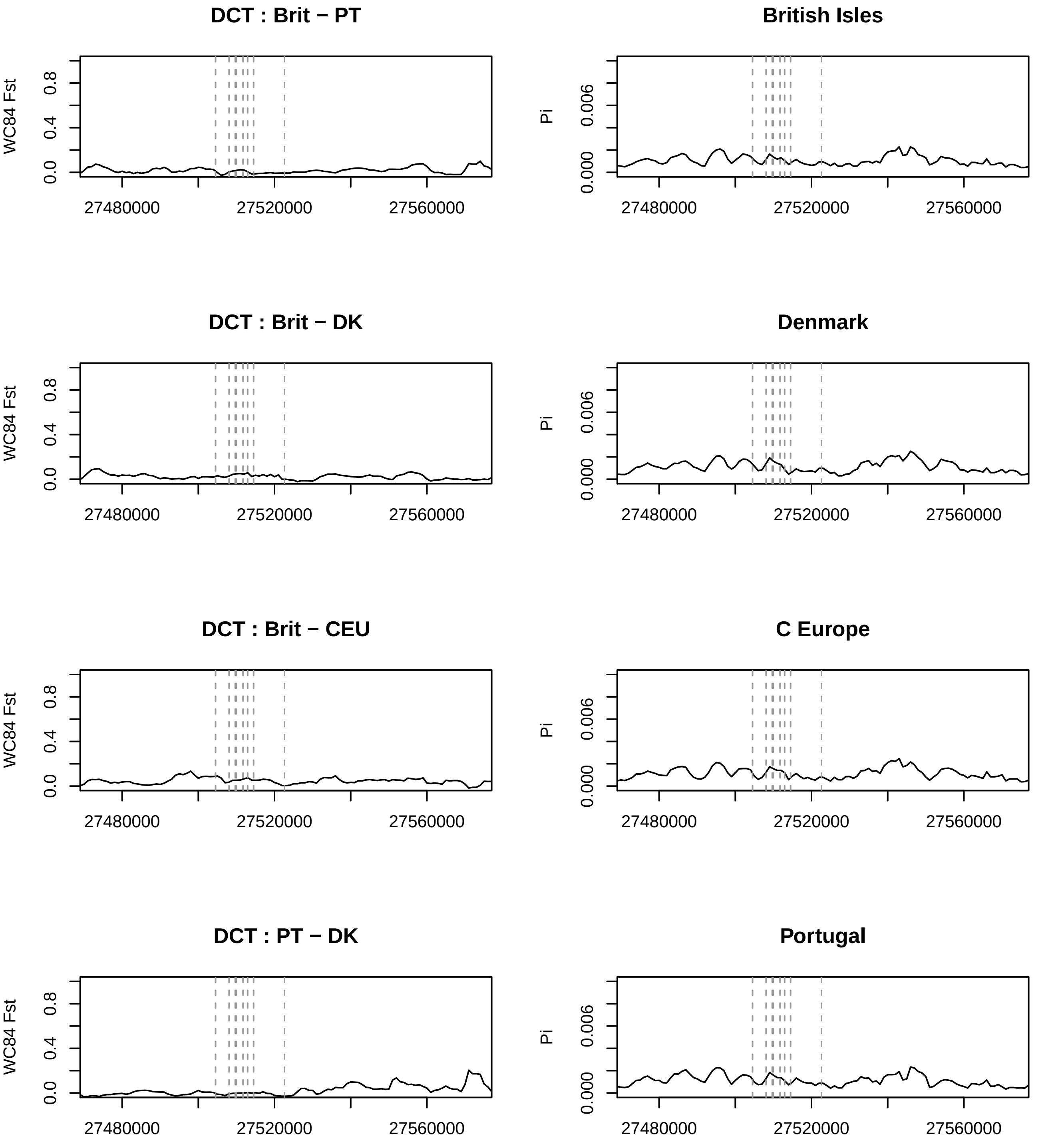
DCT : Brit − PT
British Isles
DCT : Brit − DK
Denmark
DCT : Brit − CEU
C Europe
DCT : PT − DK
Portugal
WC84 Fst
Pi
0.0
0.4
0.8
0.000
0.006
27480000
27520000
27560000

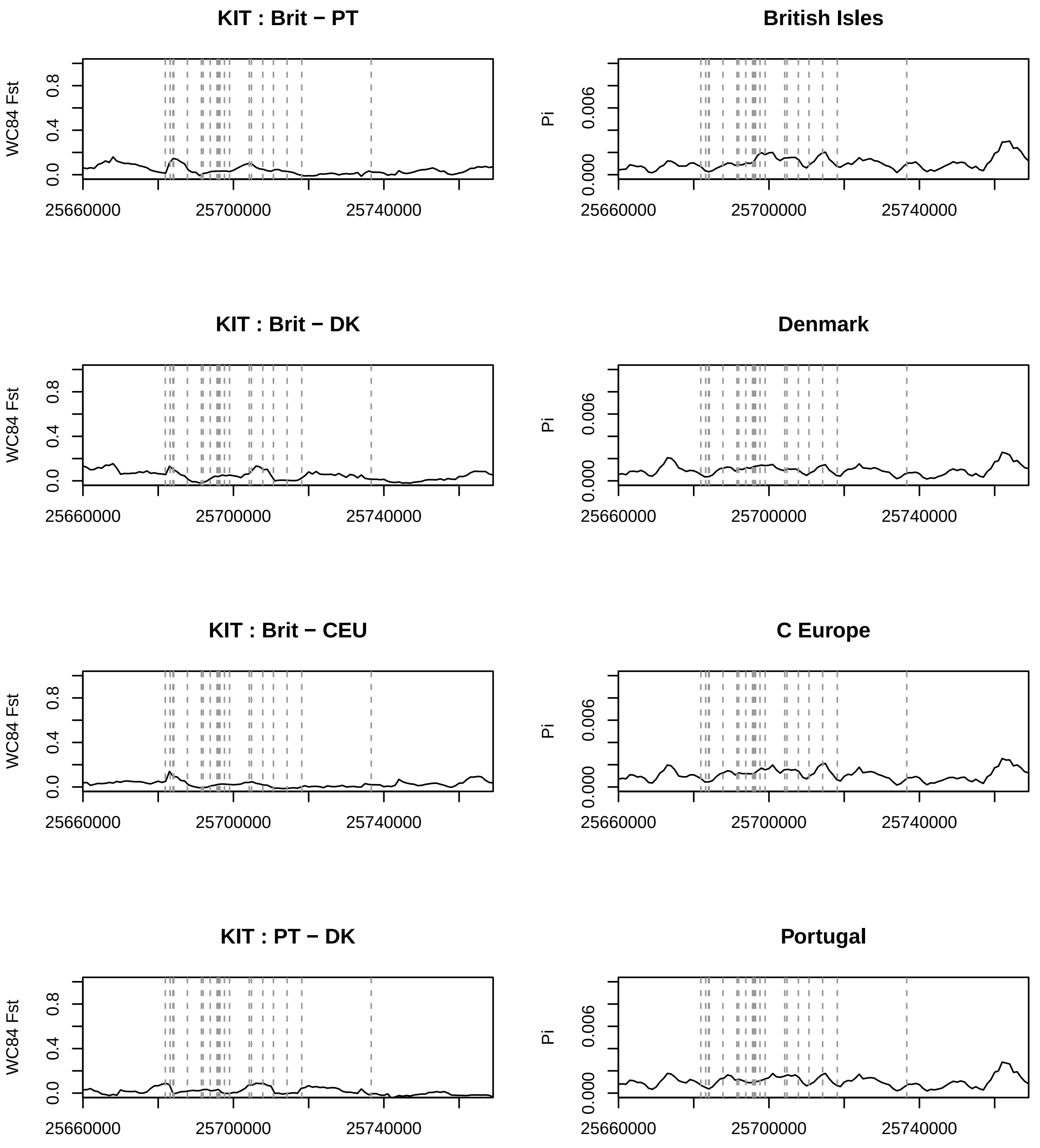
KIT : Brit − PT
British Isles
KIT : Brit − DK
Denmark
KIT : Brit − CEU
C Europe
KIT : PT − DK
Portugal
WC84 Fst
Pi
0.0
0.4
0.8
0.000
0.006
25660000
25700000
25740000

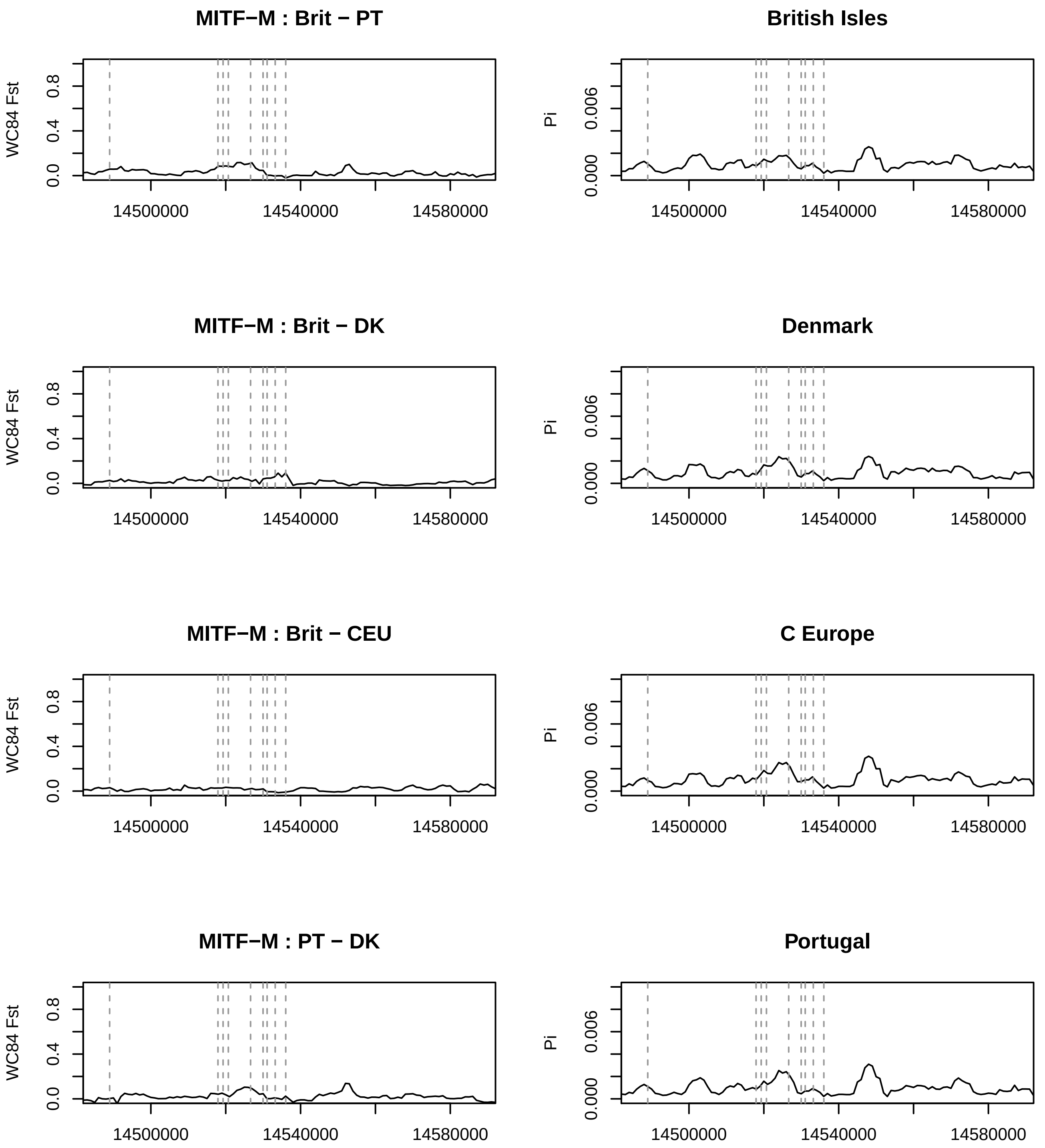
MITF−M : Brit − PT
British Isles
MITF−M : Brit − DK
Denmark
MITF−M : Brit − CEU
C Europe
MITF−M : PT − DK
Portugal
WC84 Fst
Pi
0.0
0.4
0.8
0.000
0.006
14500000
14540000
14580000

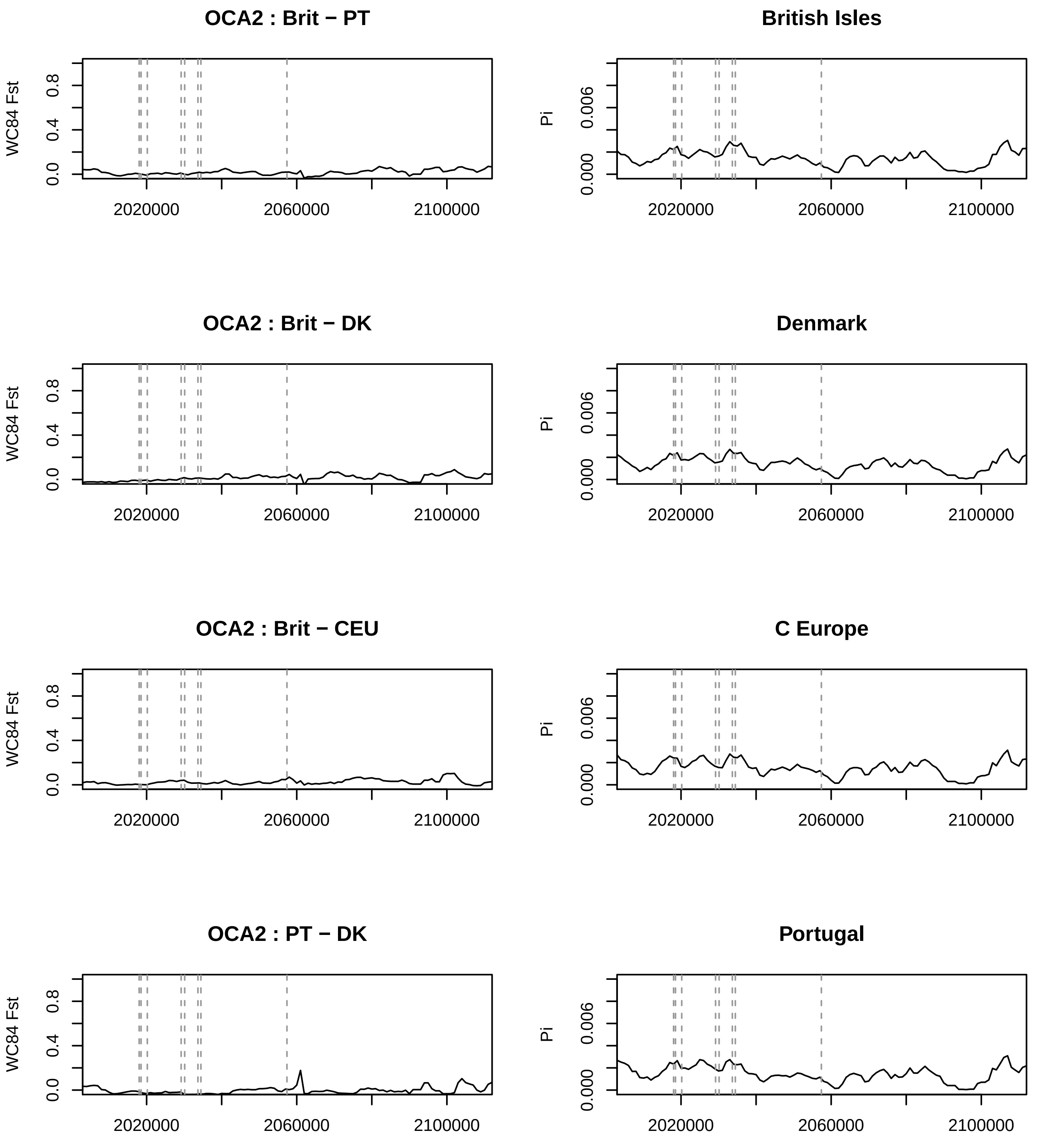

OCA2 : Brit − PT
British Isles
OCA2 : Brit − DK
Denmark
OCA2 : Brit − CEU
C Europe
OCA2 : PT − DK
Portugal
WC84 Fst
Pi
0.0
0.4
0.8
0.000
0.006
2020000
2060000
2100000

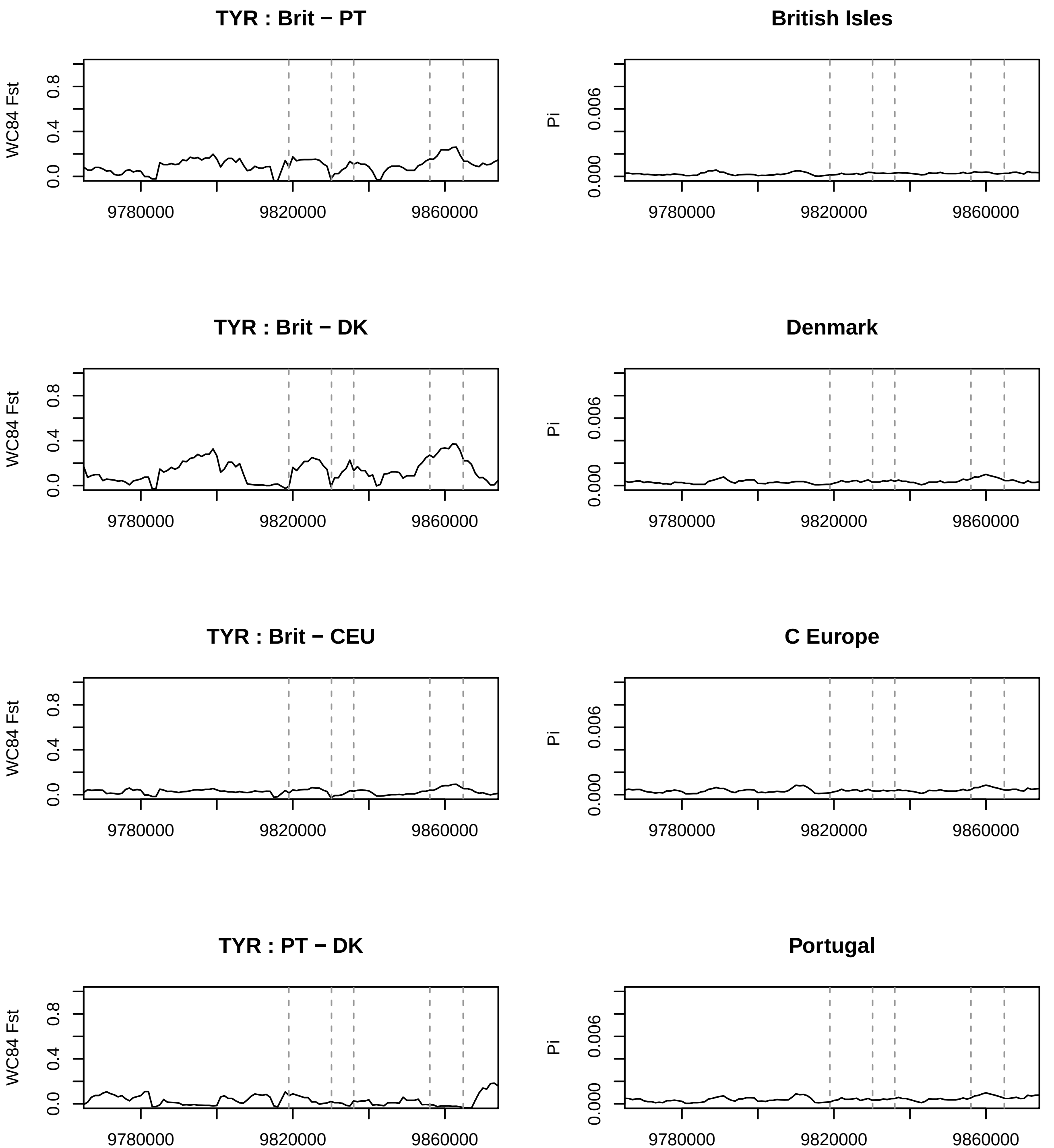
TYR : Brit − PT
British Isles
TYR : Brit − DK
Denmark
TYR : Brit − CEU
C Europe
TYR : PT − DK
Portugal
WC84 Fst
Pi
0.0
0.4
0.8
0.000
0.006
9780000
9820000
9860000

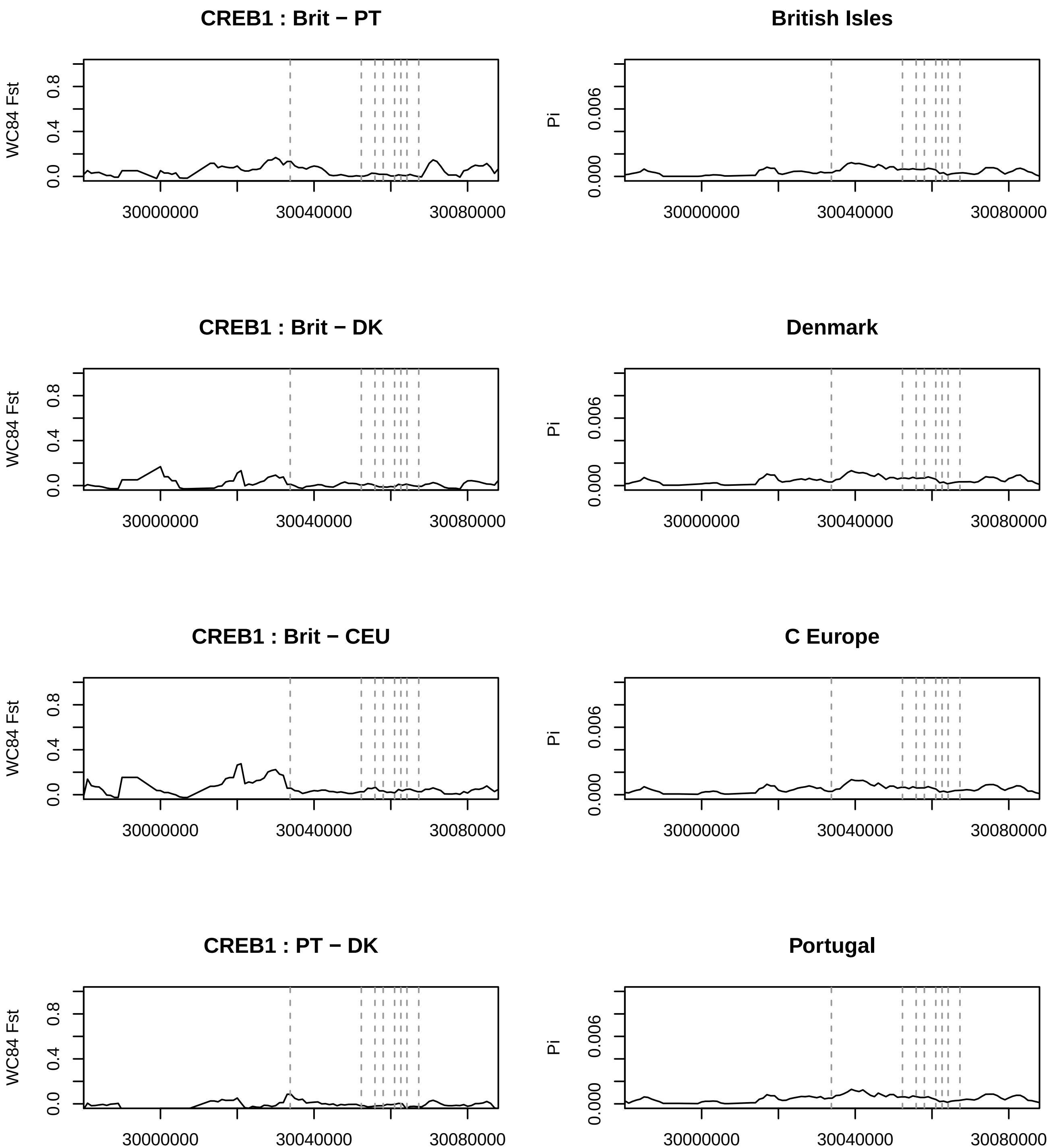
CREB1 : Brit − PT
British Isles
CREB1 : Brit − DK
Denmark
CREB1 : Brit − CEU
C Europe
CREB1 : PT − DK
Portugal
WC84 Fst
Pi
0.0
0.4
0.8
0.000
0.006
30000000
30040000
30080000

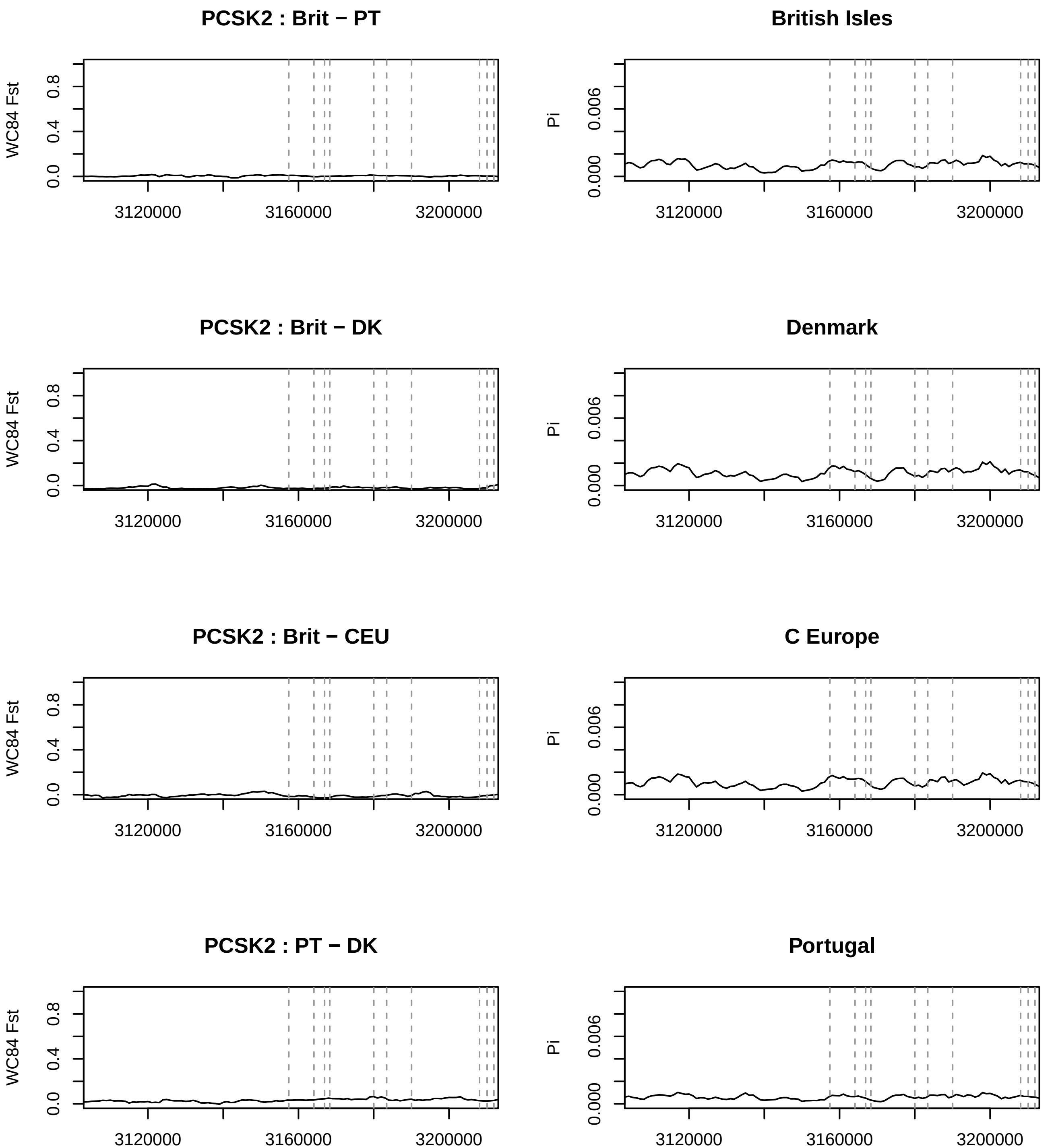
PCSK2 : Brit − PT
British Isles
PCSK2 : Brit − DK
Denmark
PCSK2 : Brit − CEU
C Europe
PCSK2 : PT − DK
Portugal
WC84 Fst
Pi
0.0
0.4
0.8
0.000
0.006
3120000
3160000
3200000

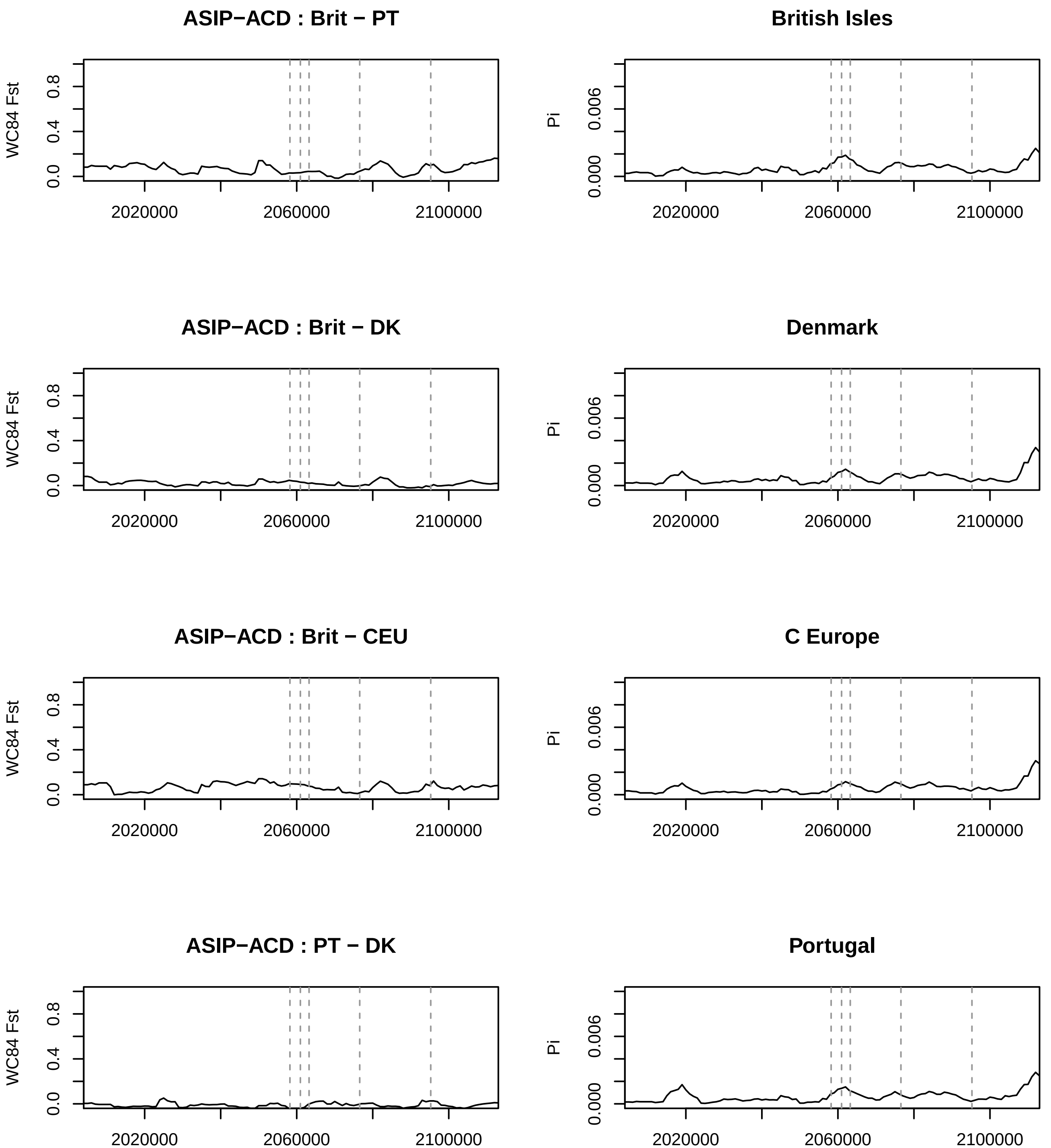
ASIP−ACD : Brit − PT
British Isles
ASIP−ACD : Brit − DK
Denmark
ASIP−ACD : Brit − CEU
C Europe
ASIP−ACD : PT − DK
Portugal
WC84 Fst
Pi
0.0
0.4
0.8
0.000
0.006
2020000
2060000
2100000

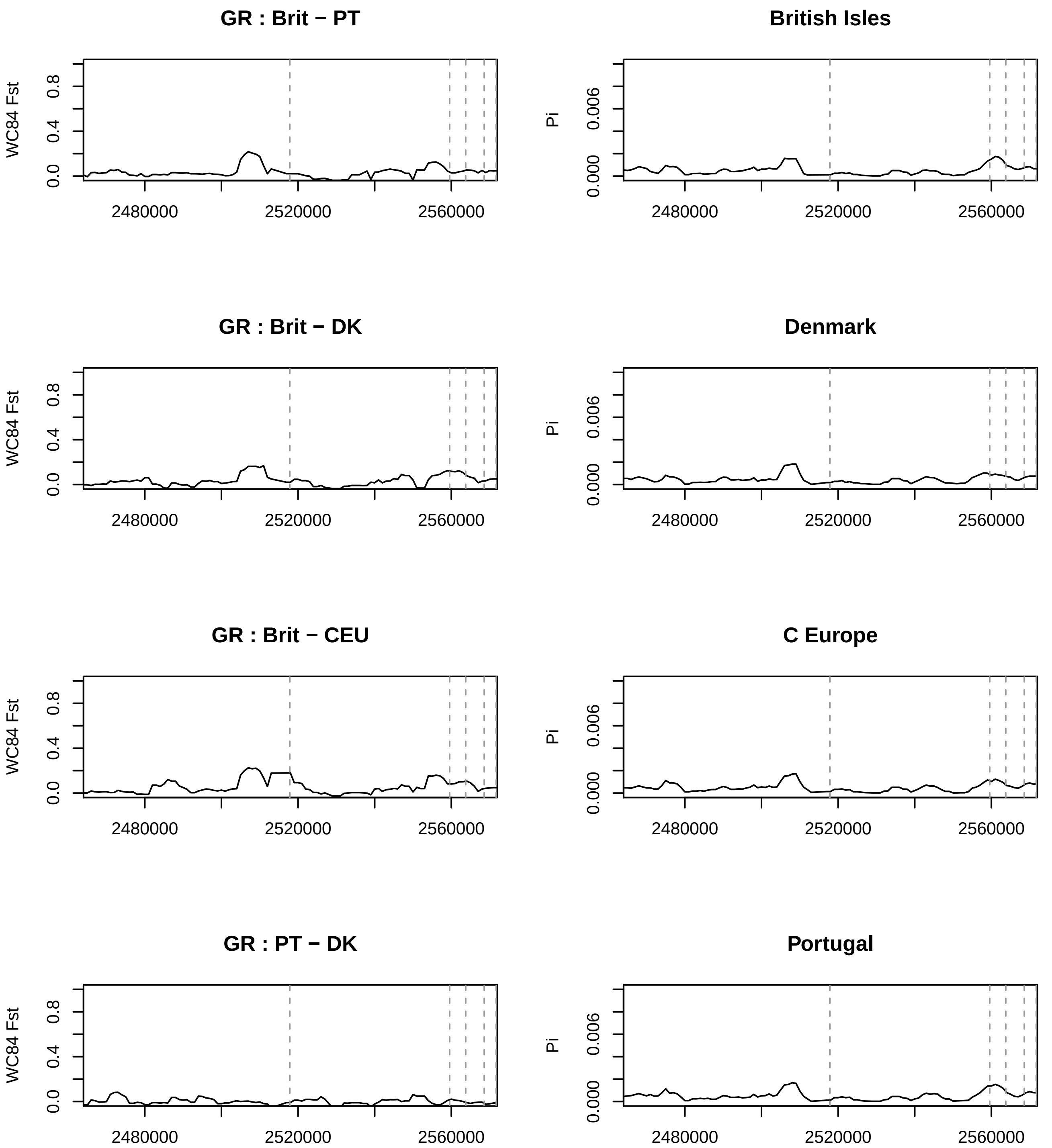
GR : Brit − PT
British Isles
GR : Brit − DK
Denmark
GR : Brit − CEU
C Europe
GR : PT − DK
Portugal
WC84 Fst
Pi
0.0
0.4
0.8
0.000
0.006
2480000
2520000
2560000

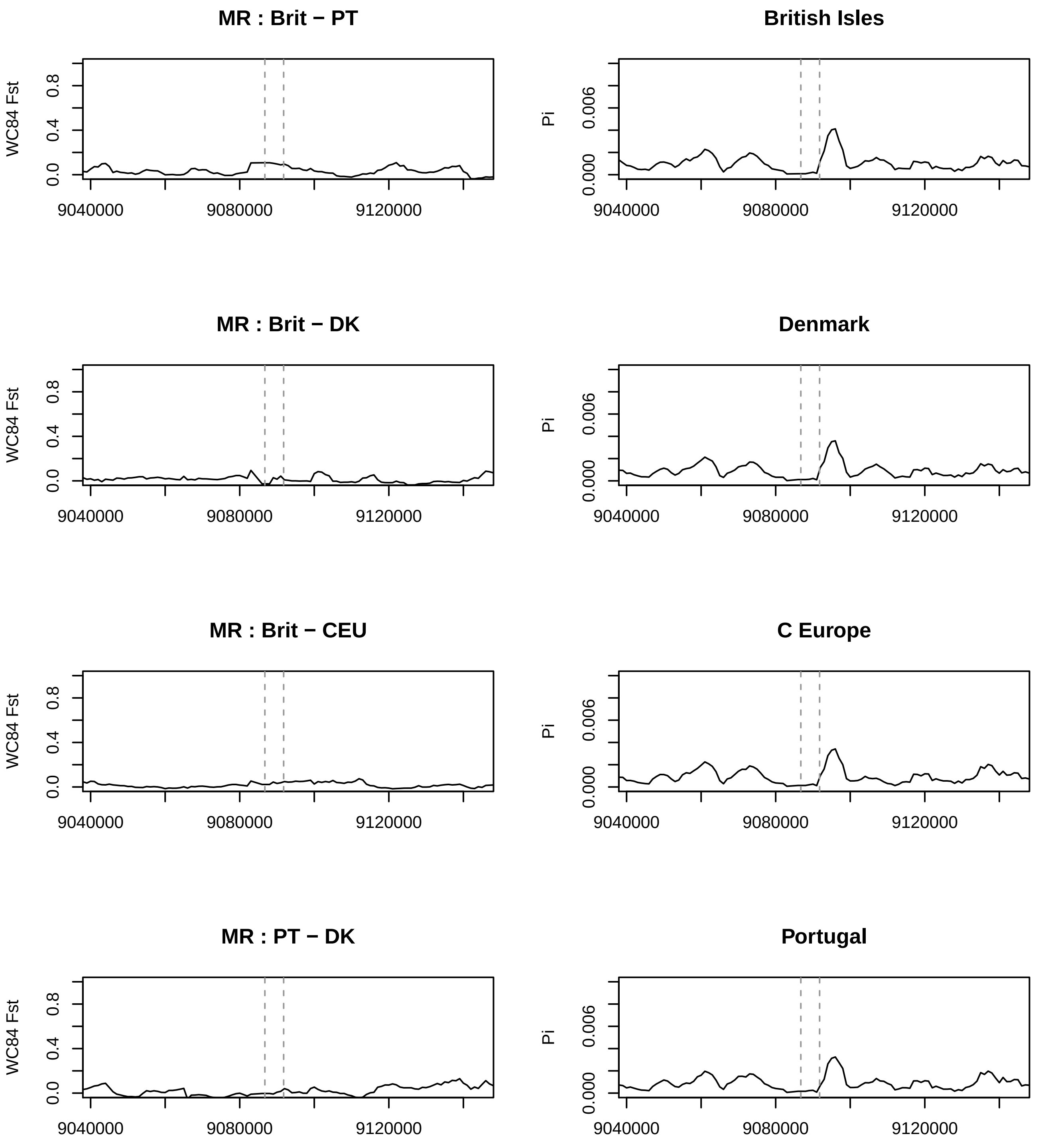
MR : Brit − PT
WC84 Fst
0.0
0.4
0.8
9040000
9080000
9120000
British Isles
Pi
0.000
0.006
MR : Brit − DK
Denmark
MR : Brit − CEU
C Europe
MR : PT − DK
Portugal

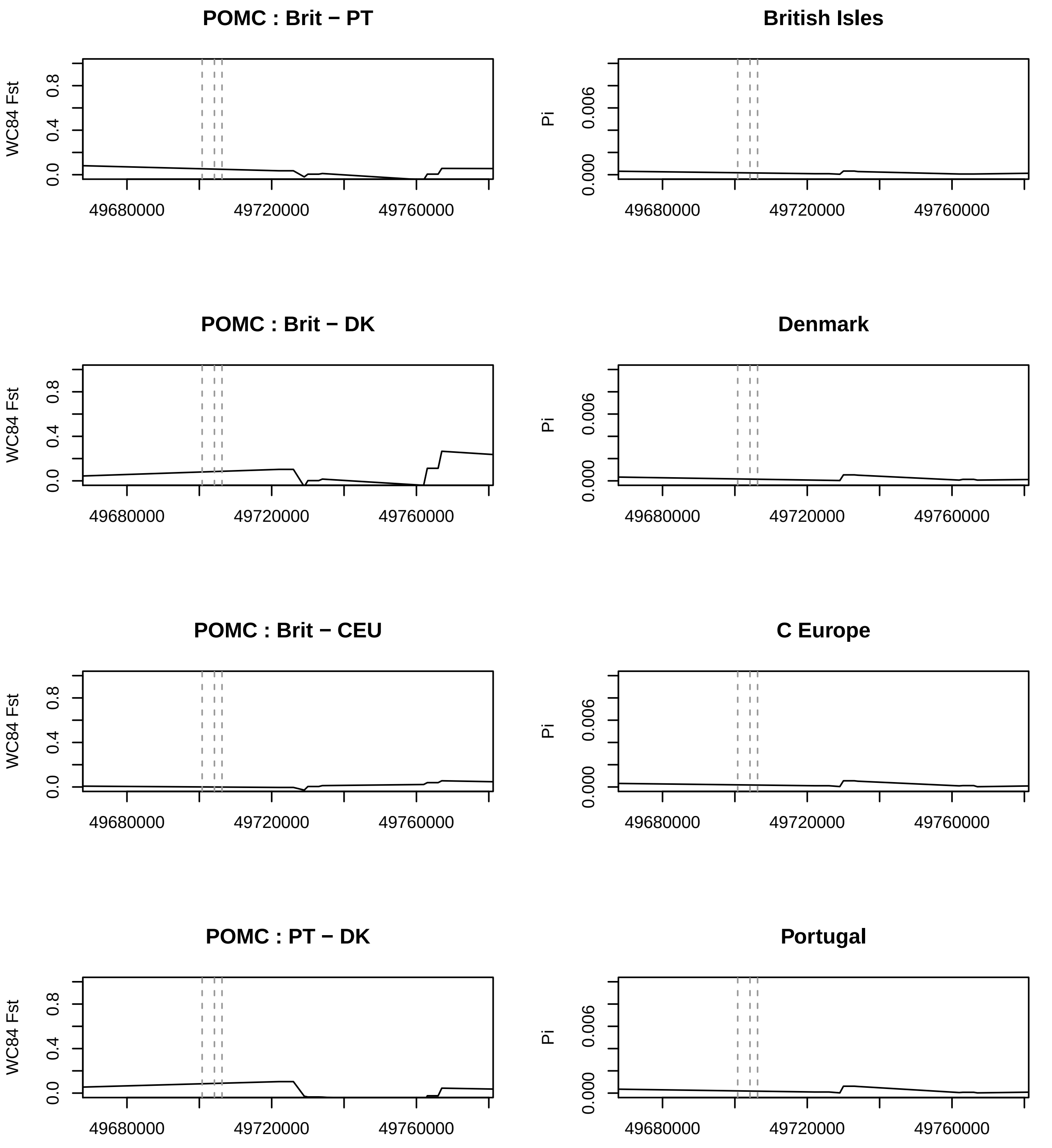
POMC : Brit − PT
WC84 Fst
0.0
0.4
0.8
49680000
49720000
49760000
British Isles
Pi
0.000
0.006
POMC : Brit − DK
Denmark
POMC : Brit − CEU
C Europe
POMC : PT − DK
Portugal

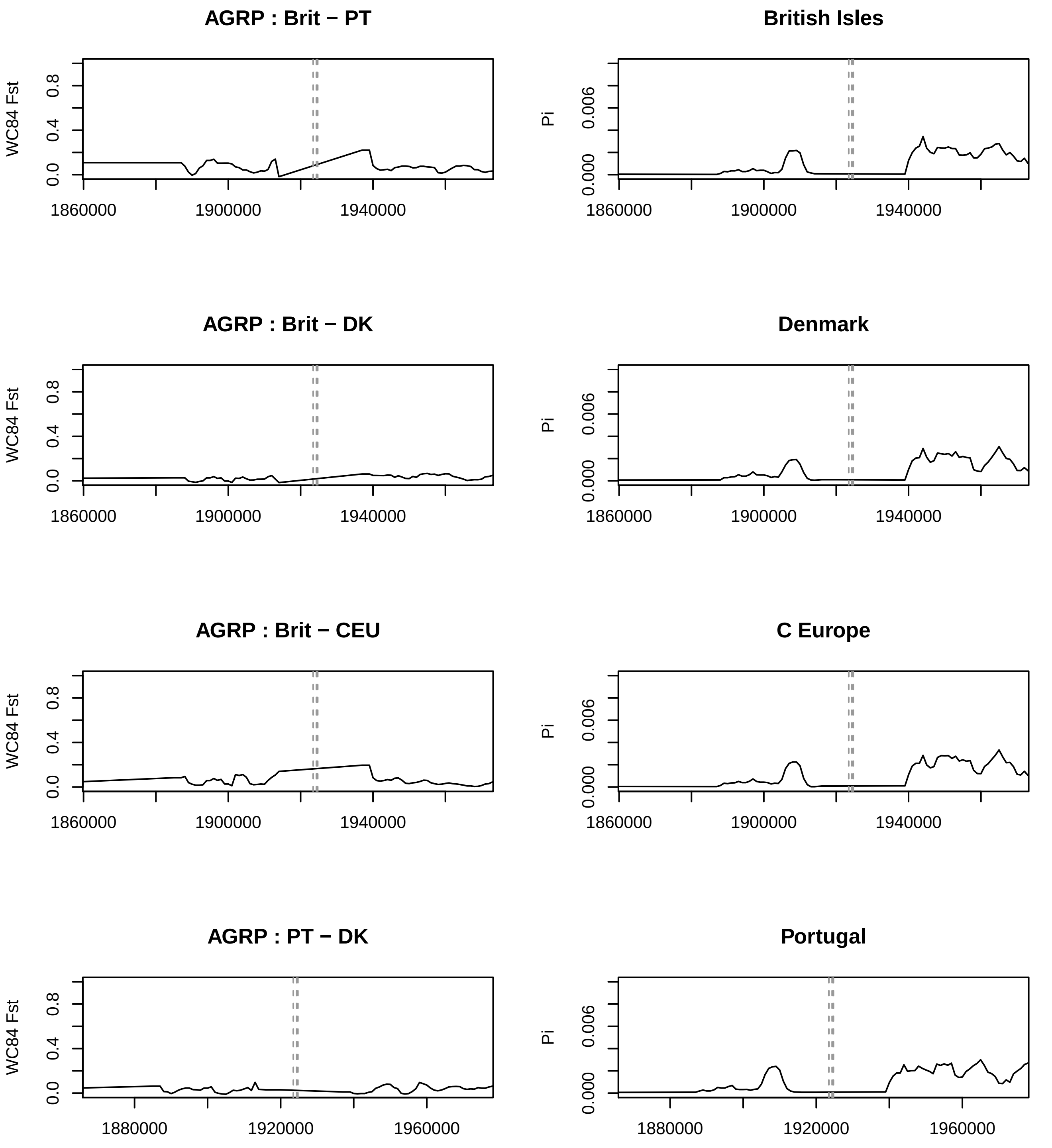
AGRP : Brit − PT
British Isles
AGRP : Brit − DK
Denmark
AGRP : Brit − CEU
C Europe
AGRP : PT − DK
Portugal
WC84 Fst
Pi
0.0
0.4
0.8
0.000
0.006
1860000
1900000
1940000
1880000
1920000
1960000

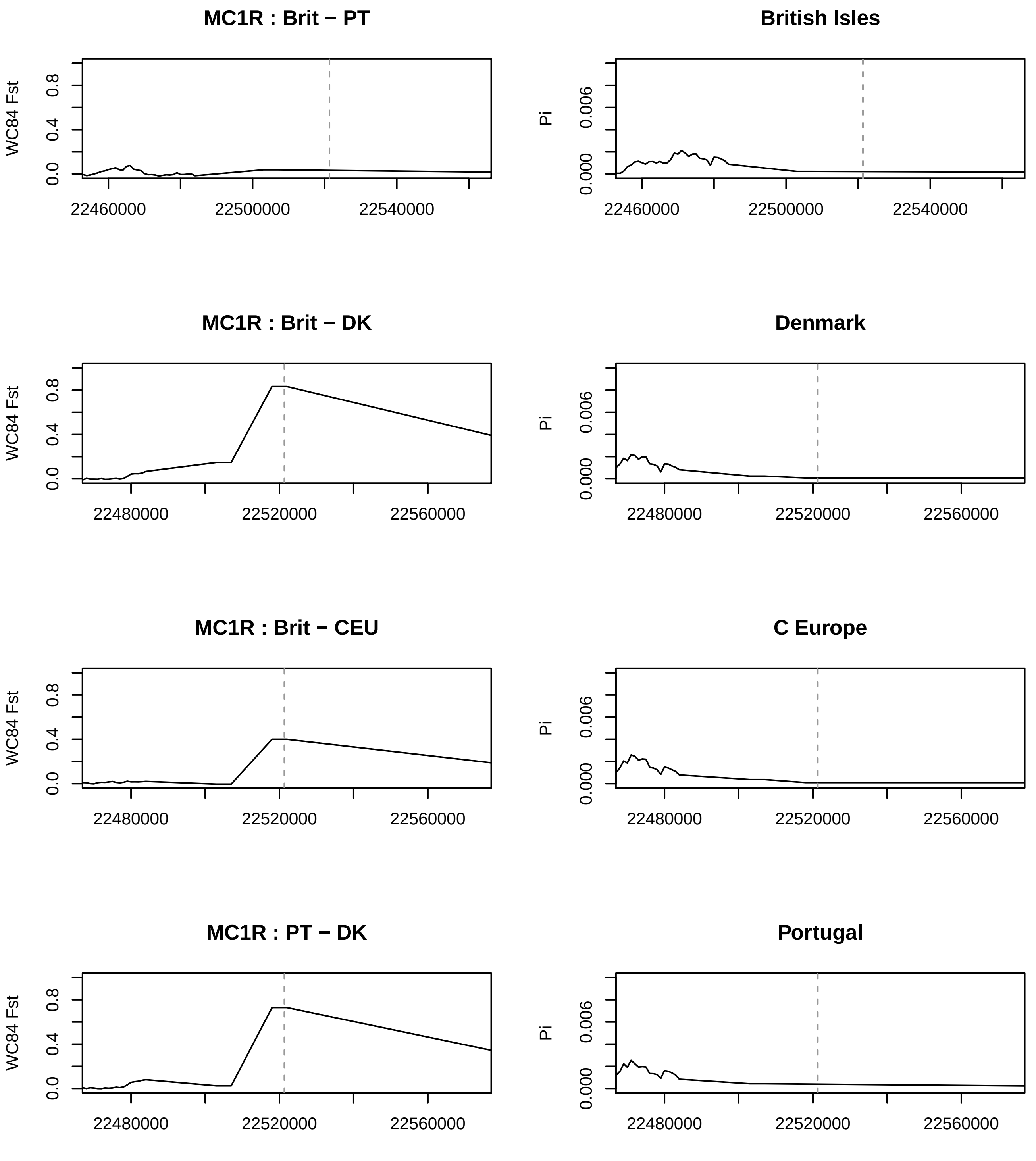

MC1R : Brit − PT
British Isles
MC1R : Brit − DK
Denmark
MC1R : Brit − CEU
C Europe
MC1R : PT − DK
Portugal
WC84 Fst
Pi
0.0
0.4
0.8
0.000
0.006
22460000
22500000
22540000
22480000
22520000
22560000

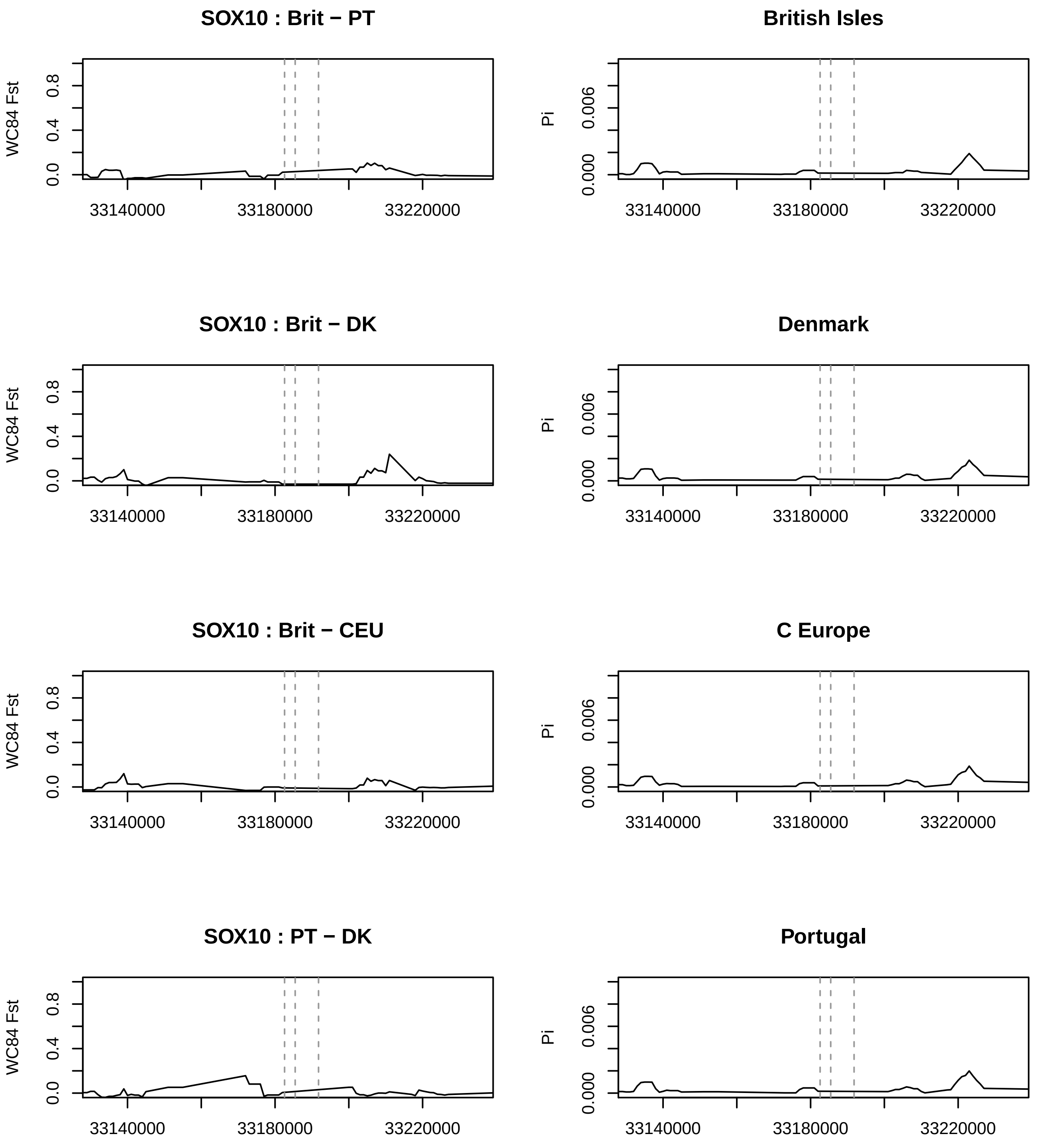
SOX10 : Brit − PT
WC84 Fst
0.0
0.4
0.8
33140000
33180000
33220000
British Isles
Pi
0.000
0.006
33140000
33180000
33220000
SOX10 : Brit − DK
WC84 Fst
0.0
0.4
0.8
33140000
33180000
33220000
Denmark
Pi
0.000
0.006
33140000
33180000
33220000
SOX10 : Brit − CEU
WC84 Fst
0.0
0.4
0.8
33140000
33180000
33220000
C Europe
Pi
0.000
0.006
33140000
33180000
33220000
SOX10 : PT − DK
WC84 Fst
0.0
0.4
0.8
33140000
33180000
33220000
Portugal
Pi
0.000
0.006
33140000
33180000
33220000

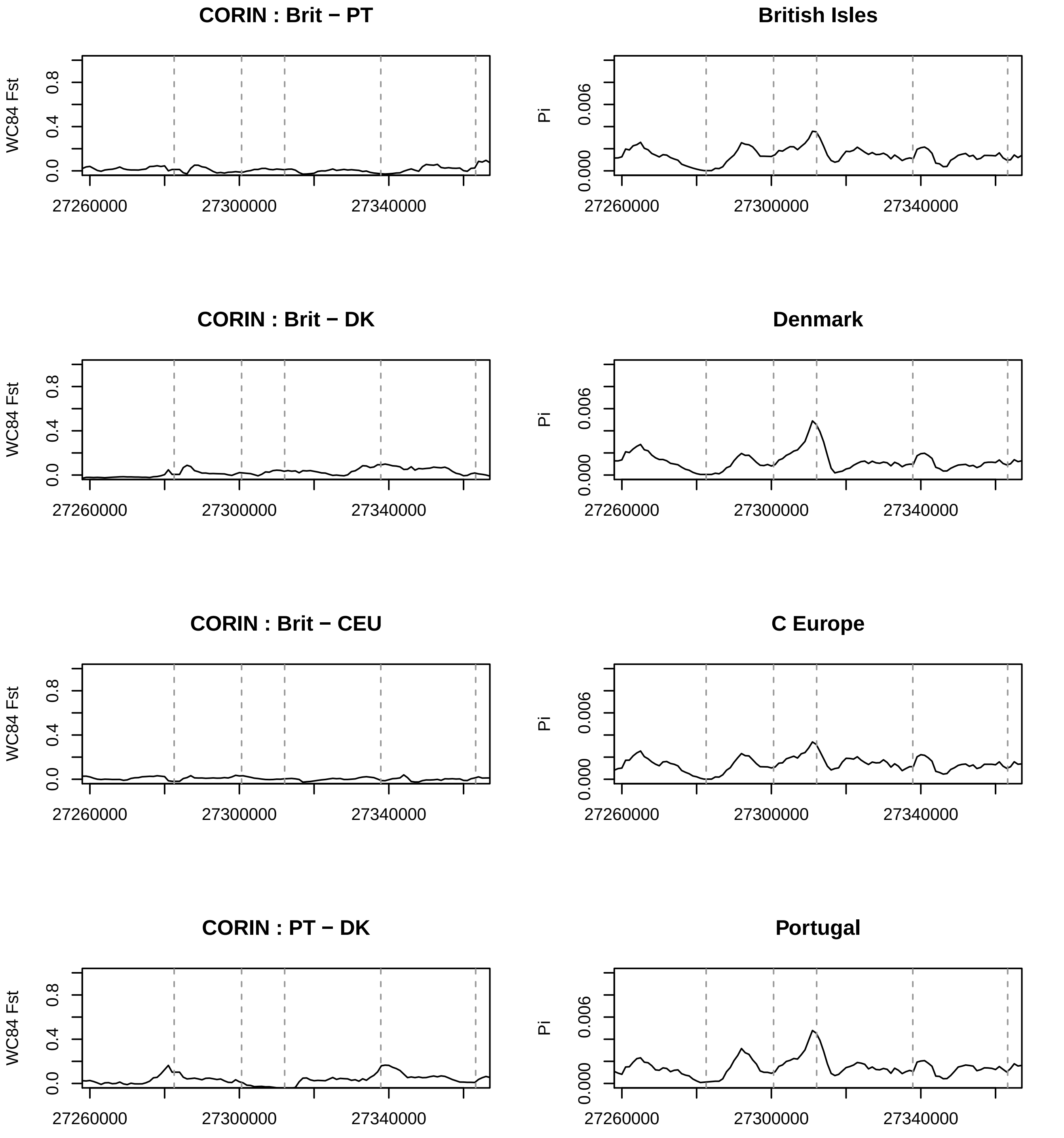
CORIN : Brit − PT
British Isles
CORIN : Brit − DK
Denmark
CORIN : Brit − CEU
C Europe
CORIN : PT − DK
Portugal
WC84 Fst
Pi
0.0
0.4
0.8
0.000
0.006
27260000
27300000
27340000

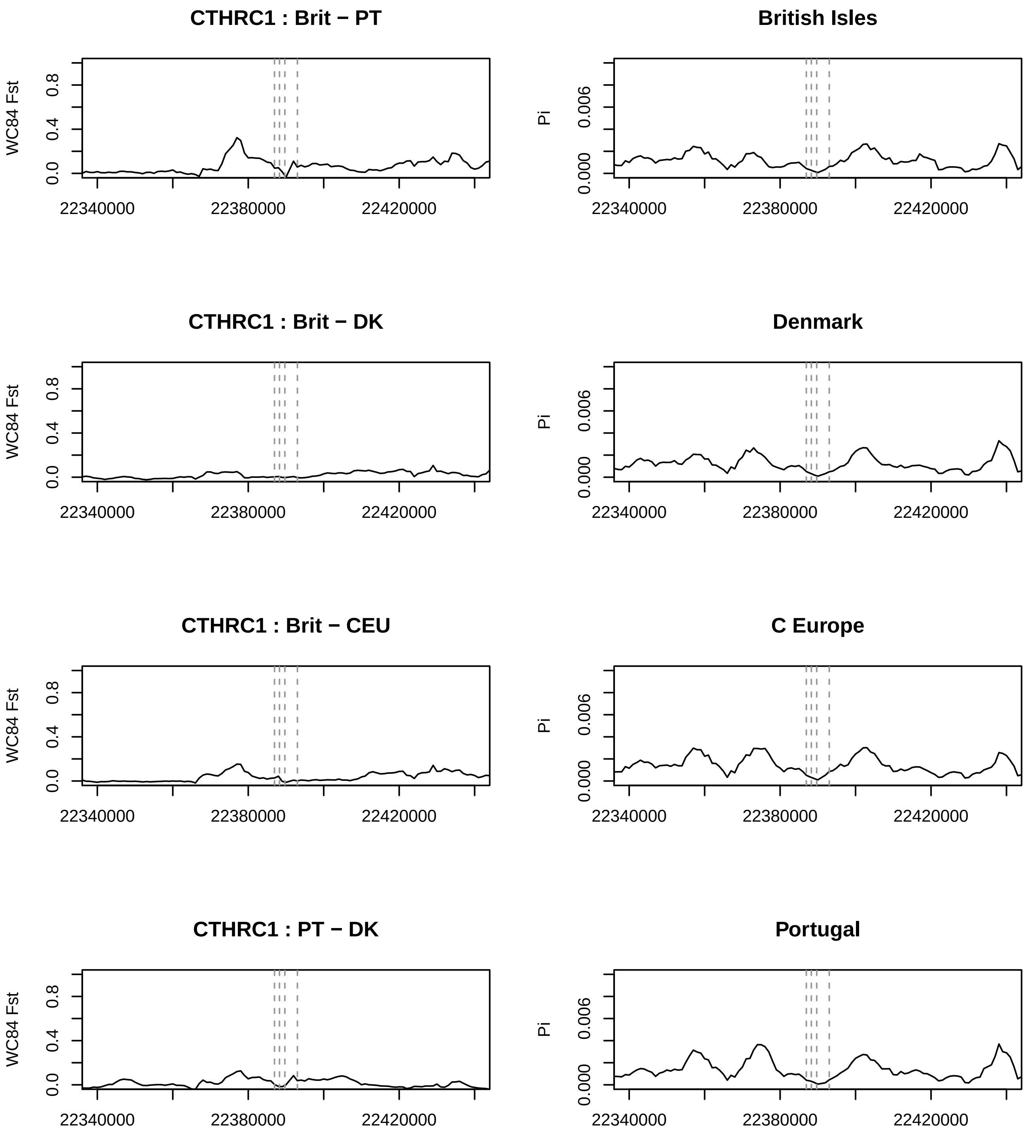
CTHRC1 : Brit − PT
British Isles
CTHRC1 : Brit − DK
Denmark
CTHRC1 : Brit − CEU
C Europe
CTHRC1 : PT − DK
Portugal
WC84 Fst
Pi
0.0
0.4
0.8
0.000
0.006
22340000
22380000
22420000

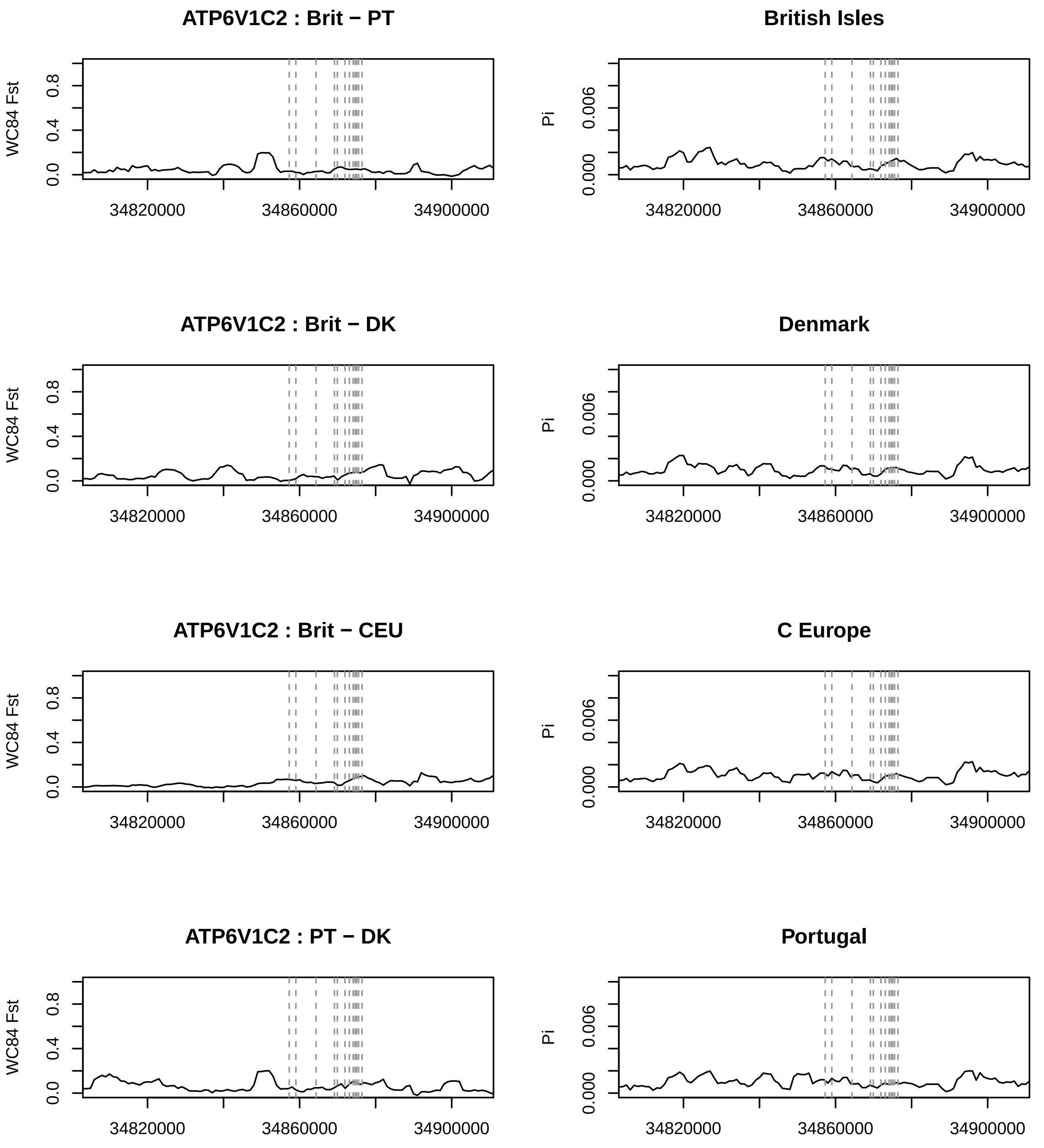

ATP6V1C2 : Brit − PT
British Isles
ATP6V1C2 : Brit − DK
Denmark
ATP6V1C2 : Brit − CEU
C Europe
ATP6V1C2 : PT − DK
Portugal
WC84 Fst
Pi
0.0
0.4
0.8
0.000
0.006
34820000
34860000
34900000

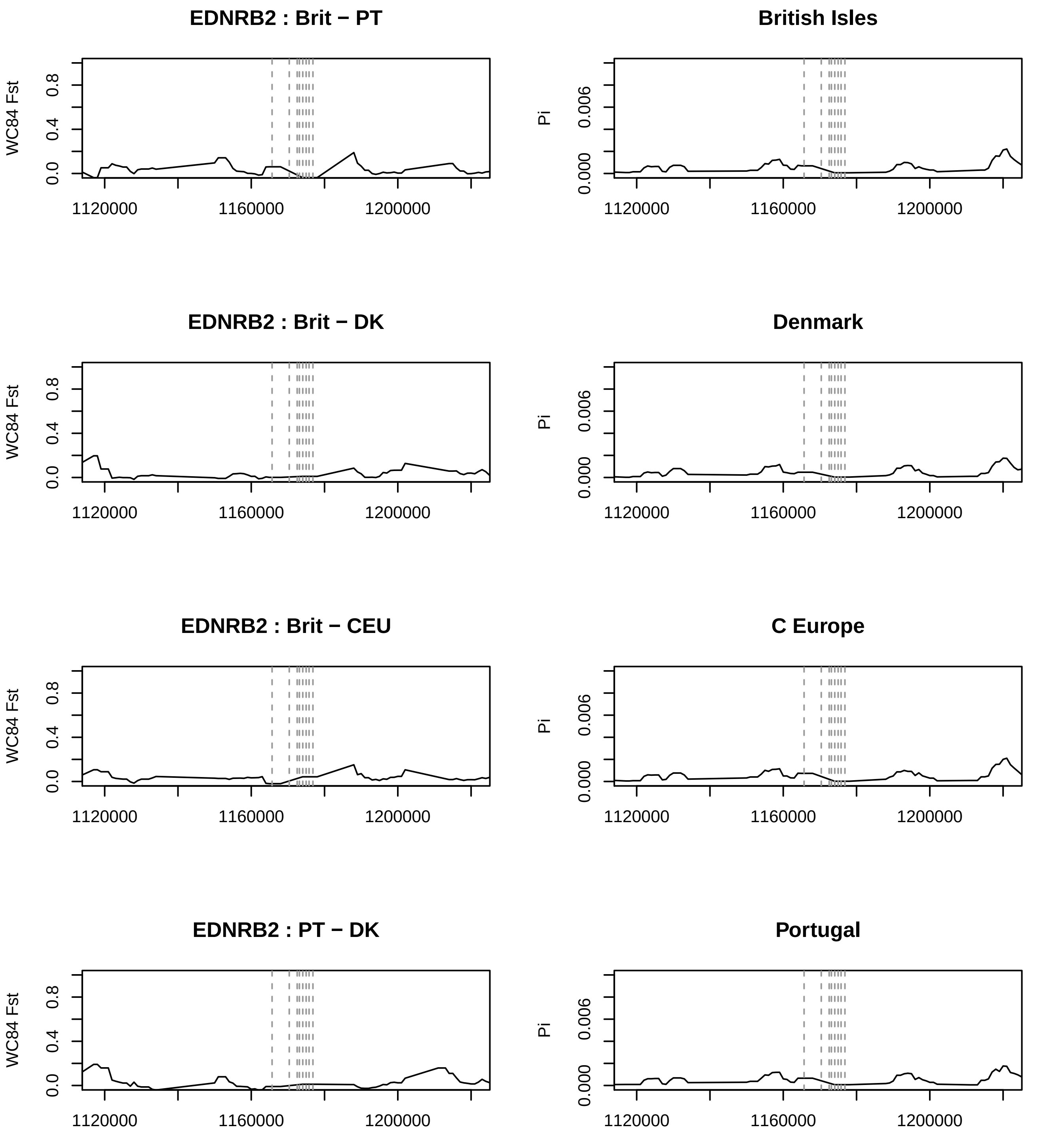
EDNRB2 : Brit − PT
British Isles
EDNRB2 : Brit − DK
Denmark
EDNRB2 : Brit − CEU
C Europe
EDNRB2 : PT − DK
Portugal
WC84 Fst
Pi
0.0
0.4
0.8
0.000
0.006
1120000
1160000
1200000

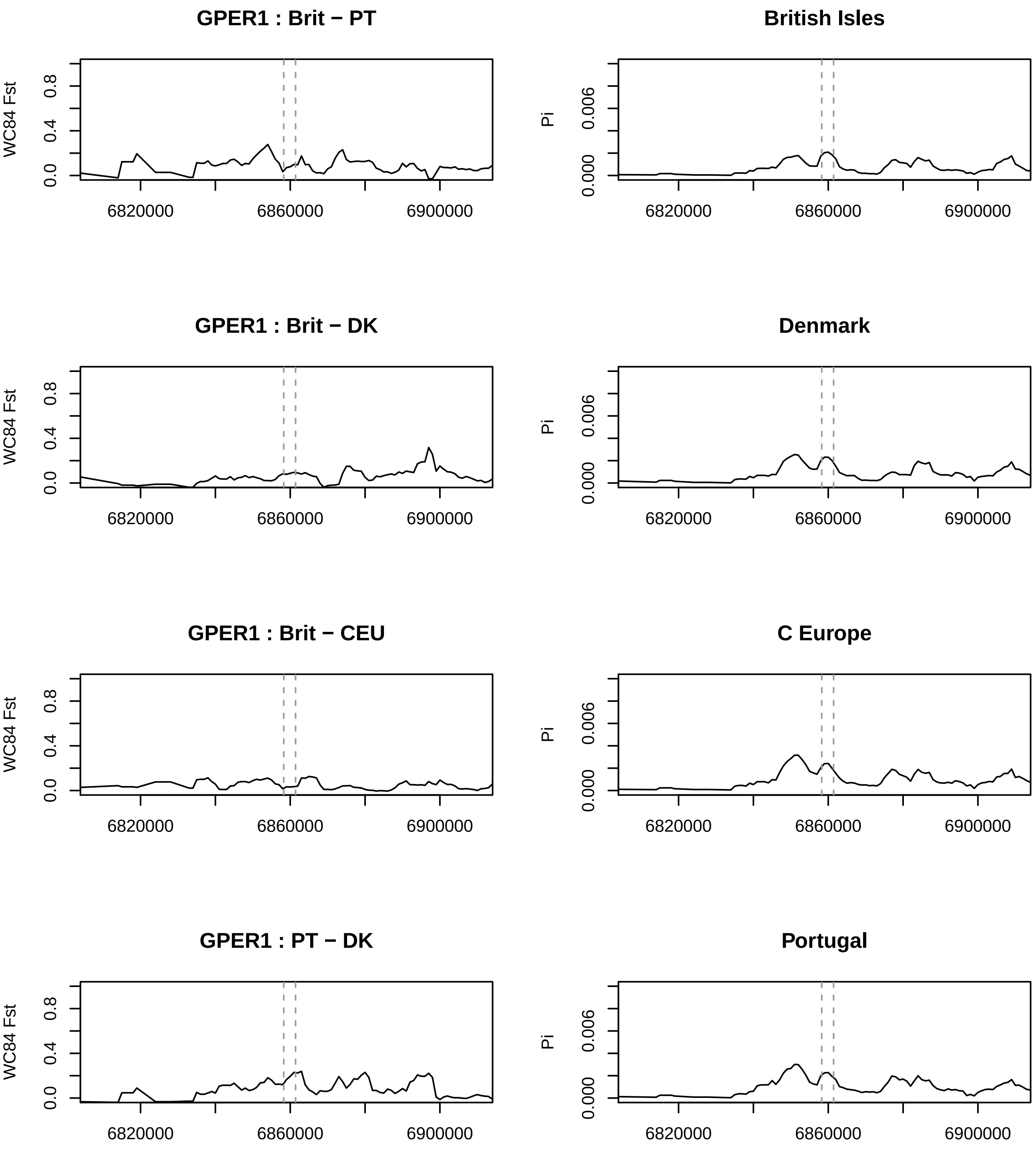
GPER1 : Brit − PT
British Isles
GPER1 : Brit − DK
Denmark
GPER1 : Brit − CEU
C Europe
GPER1 : PT − DK
Portugal
WC84 Fst
Pi
0.0
0.4
0.8
0.000
0.006
6820000
6860000
6900000

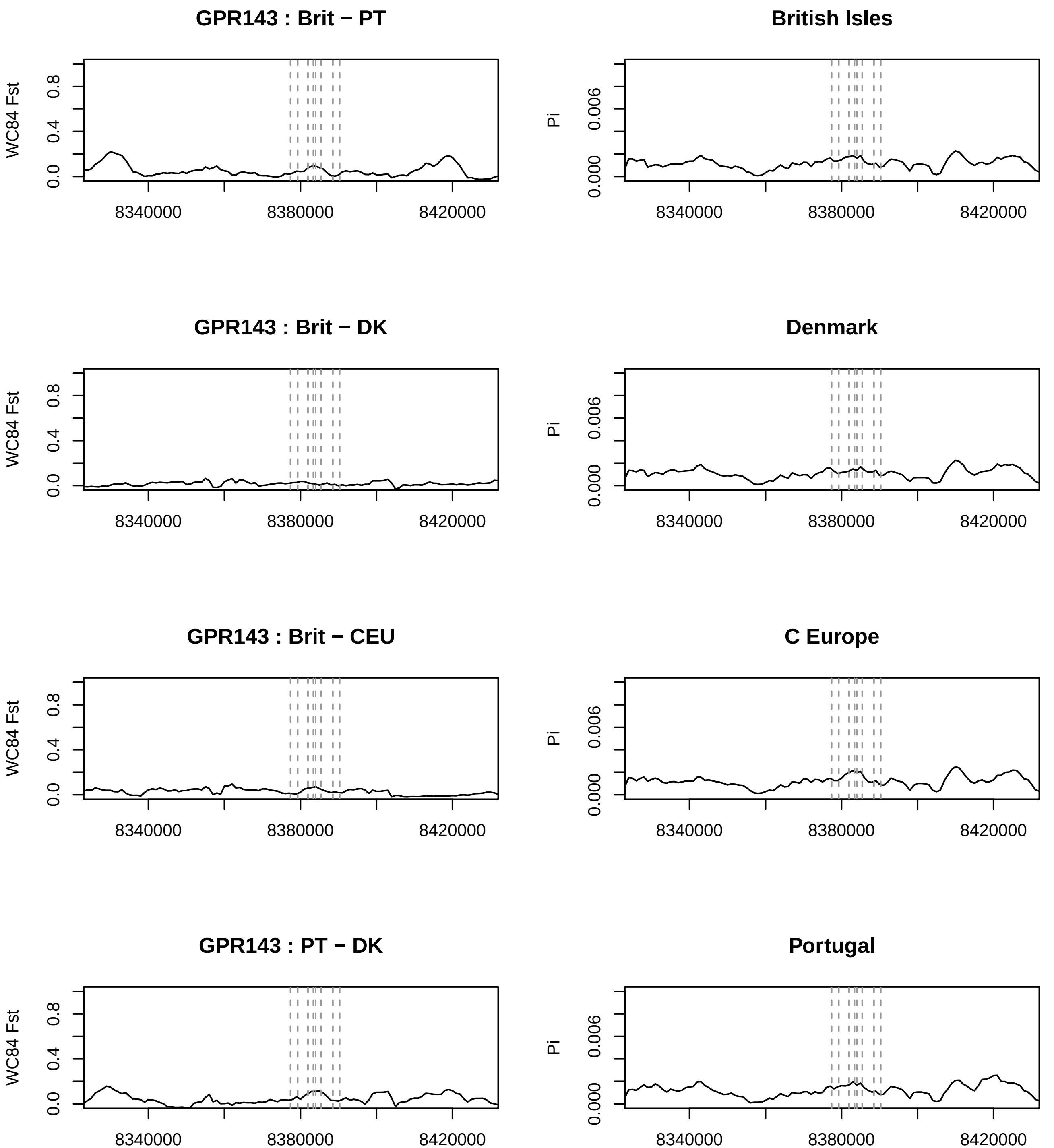
GPR143 : Brit − PT
British Isles
GPR143 : Brit − DK
Denmark
GPR143 : Brit − CEU
C Europe
GPR143 : PT − DK
Portugal
WC84 Fst
Pi
0.0
0.4
0.8
0.000
0.006
8340000
8380000
8420000

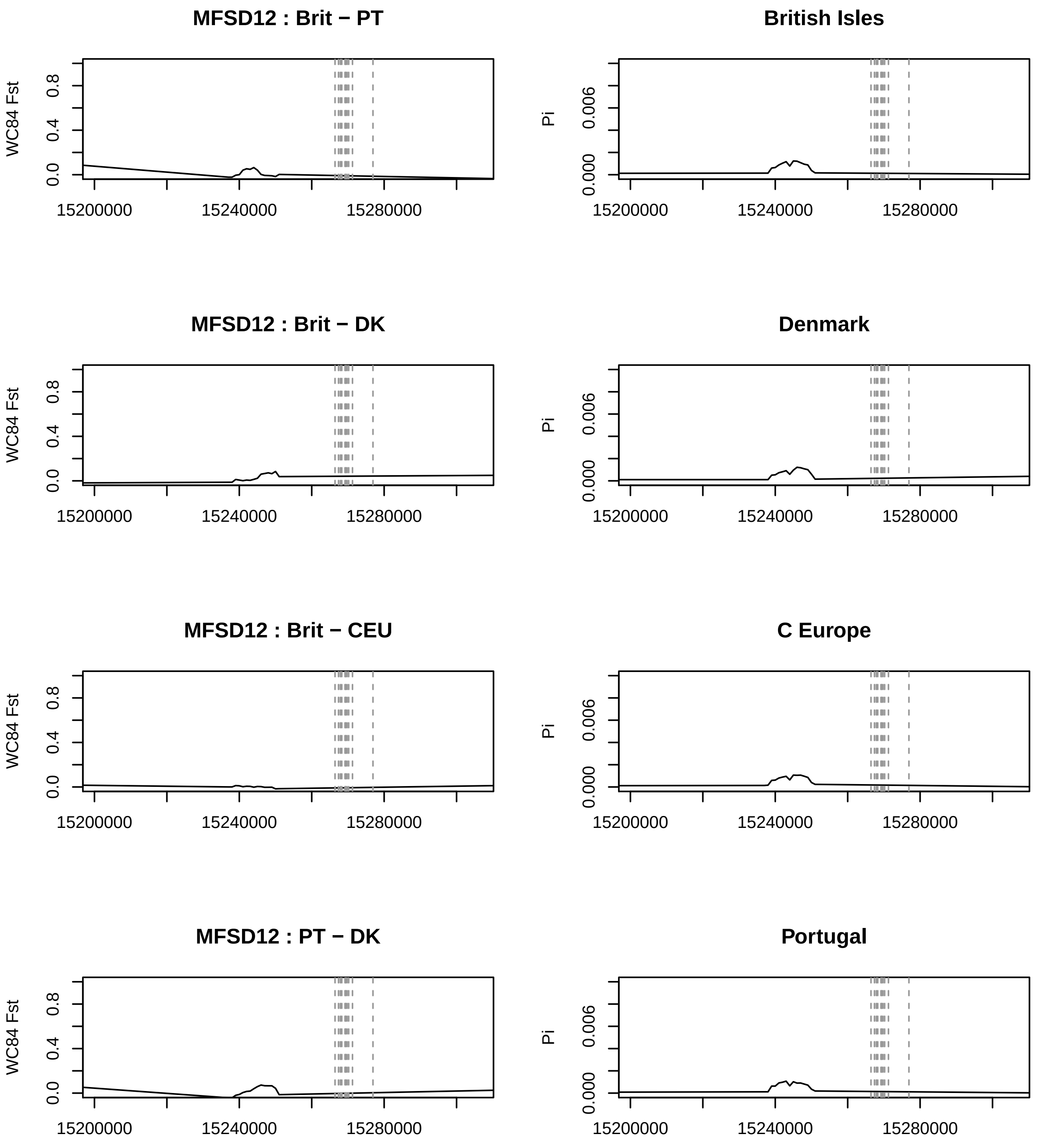

MFSD12 : Brit − PT
British Isles
MFSD12 : Brit − DK
Denmark
MFSD12 : Brit − CEU
C Europe
MFSD12 : PT − DK
Portugal
WC84 Fst
Pi
0.0
0.4
0.8
0.000
0.006
15200000
15240000
15280000

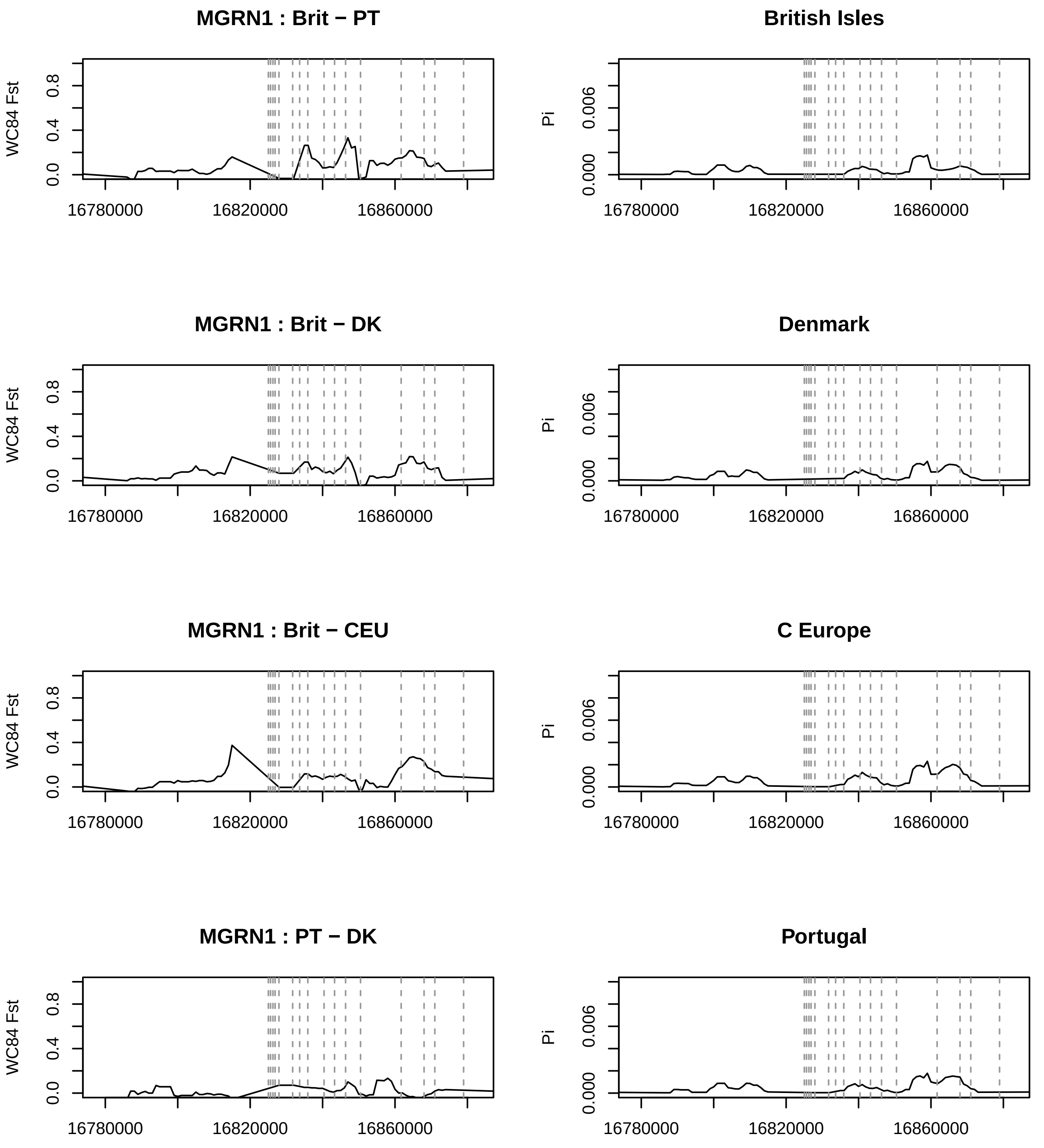
MGRN1 : Brit − PT
British Isles
MGRN1 : Brit − DK
Denmark
MGRN1 : Brit − CEU
C Europe
MGRN1 : PT − DK
Portugal
WC84 Fst
Pi
0.0
0.4
0.8
0.000
0.006
16780000
16820000
16860000

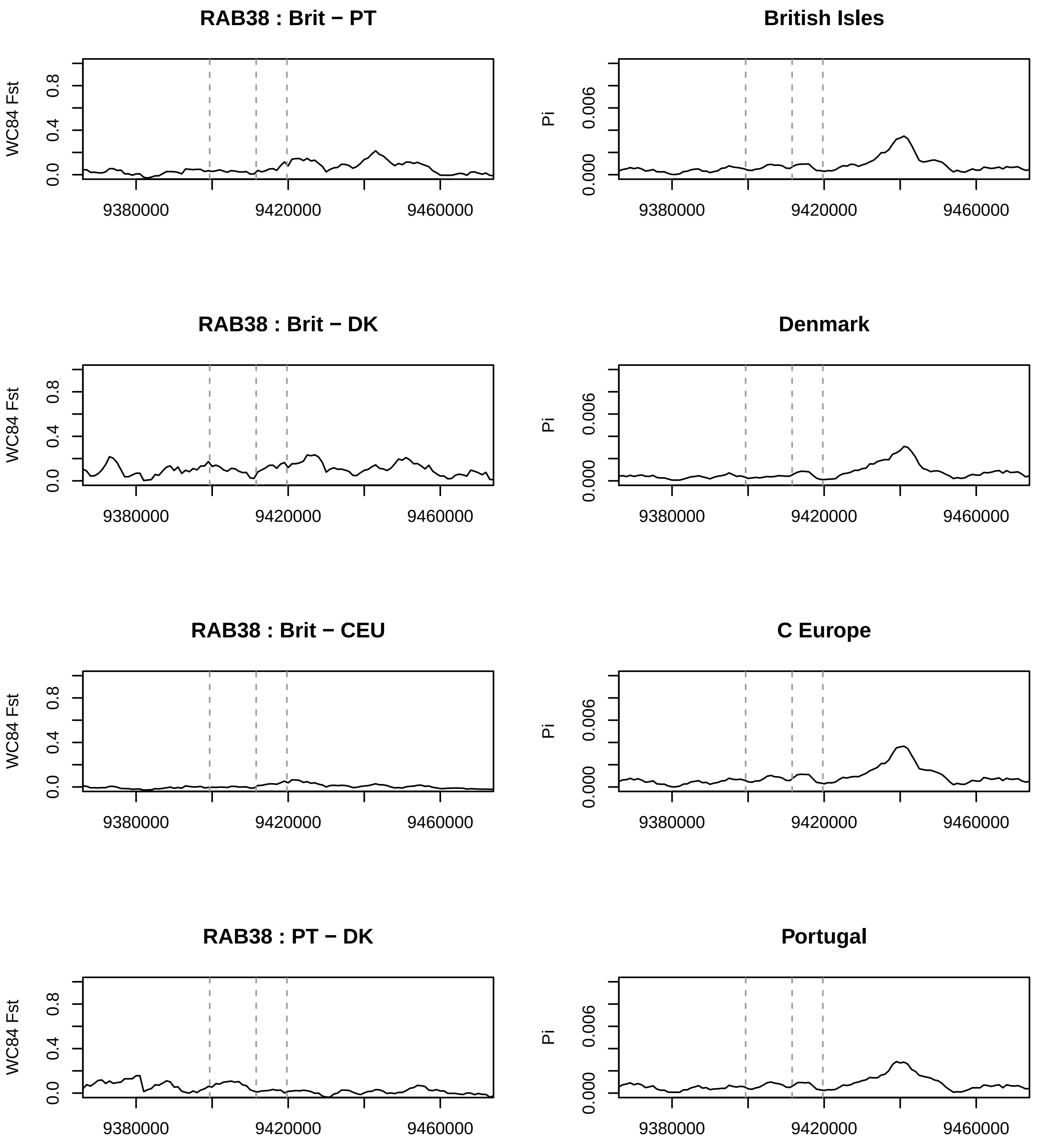
RAB38 : Brit − PT
British Isles
RAB38 : Brit − DK
Denmark
RAB38 : Brit − CEU
C Europe
RAB38 : PT − DK
Portugal
WC84 Fst
Pi
0.0
0.4
0.8
0.000
0.006
9380000
9420000
9460000
